# *In silico* analysis of DNA re-replication across a complete genome reveals cell-to-cell heterogeneity and genome plasticity

**DOI:** 10.1101/2020.03.30.016576

**Authors:** Maria Anna Rapsomaniki, Stella Maxouri, Manuel Ramirez Garrastacho, Patroula Nathanailidou, Nickolaos Nikiforos Giakoumakis, Stavros Taraviras, John Lygeros, Zoi Lygerou

## Abstract

DNA replication is a complex and remarkably robust process: despite its inherent uncertainty, manifested through stochastic replication timing at a single-cell level, multiple control mechanisms ensure its accurate and timely completion across a population. Disruptions in these mechanisms lead to DNA re-replication, closely connected to genomic instability and oncogenesis. We present a stochastic hybrid model of DNA re-replication that accurately portrays the interplay between discrete dynamics, continuous dynamics, and uncertainty. Using experimental data on the fission yeast genome, model simulations show how different regions respond to re-replication, and permit insight into the key mechanisms affecting re-replication dynamics. Simulated and experimental population-level profiles exhibit good correlation along the genome, which is robust to model parameters, validating our approach. At a single-cell level, copy numbers of individual loci are affected by intrinsic properties of each locus, *in cis* effects from adjoining loci and *in trans* effects from distant loci. *In silico* analysis and single-cell imaging reveal that cell-to-cell heterogeneity is inherent in re-replication and can lead to a plethora of genotypic variations. Our thorough *in silico* analysis of DNA re-replication across a complete genome reveals that heterogeneity at the single cell level and robustness at the population level are emerging and co-existing principles of DNA re-replication. Our results indicate that re-replication can promote genome plasticity by generating many diverse genotypes within a population, potentially offering an evolutionary advantage in cells with aberrations in replication control mechanisms.

## Introduction

DNA replication ensures the maintenance of genetic information and constitutes the basis of biological inheritance. In eukaryotes, DNA replication initiates at multiple sites across the genome, known as origins of replication, and continues bidirectionally through replication forks that move continuously until precisely two DNA copies are produced (Siddiqui et al., 2013; Symeonidou et al., 2012). DNA replication is a complex and uncertain process, as only a small fraction of all putative origins is selected to fire in each cell, resulting in an individual progression along the genome at a single-cell level (Rhind et al., 2010). Despite this high degree of stochasticity, DNA replication is also remarkably robust: it is tightly regulated in time and space by multiple control mechanisms that ensure its completion in an accurate and timely manner (Fragkos et al., 2015; Legouras et al., 2006; Symeonidou et al., 2012).

To maintain genome stability, each part of the genome must be replicated once and only once every time a cell divides. At the beginning of each cell cycle, two licensing factors, Cdt1 and Cdc6/18, load the MCM2-7 replicative helicase onto DNA, thereby licensing origins for a new round of DNA replication (Nishitani and Lygerou, 2004; Parker et al., 2017). In S-phase, the replicative helicase either becomes active and moves away from origins with the replication fork, or is removed by passive replication. Cdt1 and Cdc6/18 are strictly controlled and are inactivated as soon as replication starts, ensuring that the replicative helicase cannot load again onto origins that have been replicated, and therefore origins cannot fire a second time. Disruption of this control mechanism leads to re-firing of origins within the same cell cycle, a pathological process known as DNA re-replication (Blow and Dutta, 2005). Over-expression of the licensing factors Cdt1 and Cdc6/18 has been shown to promote re-replication from yeast to humans. In fission yeast, over-expression of Cdc18 leads to origin re-licensing within the same cell cycle, origin re-firing, and an uneven increase of DNA copy number (Nishitani and Nurse, 1995), resulting in local amplification of the genome (Kiang et al., 2010; Mickle et al., 2007). Re-replication is enhanced by concomitant expression of Cdt1 (Nishitani et al., 2000; Yanow, 2001). In mammalian cells, ectopic expression of Cdt1 alone or in combination with Cdc6 is sufficient to drive re-replication (Vaziri et al., 2003). Both Cdt1 and Cdc6 are often over-expressed in human tumors (Karakaidos et al., 2004), and have been linked to genomic instability early in the tumorigenesis process (Liontos et al., 2007), which drives oncogenesis (Champeris Tsaniras et al., 2014; Gaillard et al., 2015; Hills and Diffley, 2014; Hook et al., 2007; Nowak et al., 2002; Petropoulou, 2008).

Studies of re-replication in fission yeast have shown that, at the population level, re-replication progresses relatively evenly across the genome (Kiang et al., 2010; Mickle et al., 2007), while a small number of prominent loci are re-replicated above the mean. Common features of the central origins underlying these re-replicating “hotspots” include AT-richness, early firing in a normal S phase, and localization in large intergenic regions (Kiang et al., 2010), features which also characterize efficient origins (Heichinger et al., 2006). Re-replication and normal S phase origins largely overlap, however notable differences between specific loci suggest that re-replication dynamics differ from normal replication.

To date, a number of mathematical and computational models of DNA replication in a number of organisms have been developed (Blow and Ge, 2009; Gauthier and Bechhoefer, 2009; Gauthier et al., 2012; Gispan et al., 2017; Goldar et al., 2008; Kelly and Callegari, 2019; Lygeros et al., 2008; Al Mamun et al., 2016; de Moura et al., 2010). However, the properties and underlying mechanisms of DNA re-replication across the genome remain unknown. Motivated by this gap in the literature, in this work we present a realistic, dynamic model of DNA re-replication exploiting stochastic hybrid systems. Stochastic hybrid systems combine discrete and continuous states and stochasticity (2018) and have been successfully used to capture complex biological processes (Cinquemani et al., 2008; Kouretas et al., 2006; Lygeros et al., 2008; Rapsomaniki et al., 2015). Using as input experimentally determined origin measurements from fission yeast, the model allows the simulation of DNA re-replication genome-wide. Sensitivity analysis showed that the model is robust and consistent with experimental data genome-wide, allowing rules governing re-replication to be unveiled. *In silico* analysis combined with *in cell* validation showed that re-replication profiles at the single-cell level are characterized by a high degree of heterogeneity. Re-replication can, with varying probability, occur anywhere in the genome, and generate many diverse genotypes within a population.

## Results

### Modeling DNA re-replication across a complete genome

DNA replication initiates from hundreds of origins along the genome and results in the exact duplication of the genetic material (Fig. 1A). It is a complex process that involves a combination of discrete dynamics (associated with the switch-like activation of each origin), continuous dynamics (associated with the movement of the replication forks along the DNA strands), and stochasticity (in the time and space of origin firing). Mathematical and computational models of DNA replication have been proposed in the literature to capture these mixed dynamics (Blow and Ge, 2009; Gauthier and Bechhoefer, 2009; Gauthier et al., 2012; Goldar et al., 2008; Koutroumpas and Lygeros, 2011; Lygeros et al., 2008; de Moura et al., 2010). We developed an extension of a mathematical model of normal DNA replication (Lygeros et al., 2008) to allow origin re-firing, resulting in a stochastic hybrid model of DNA re-replication.

**Figure 1.**
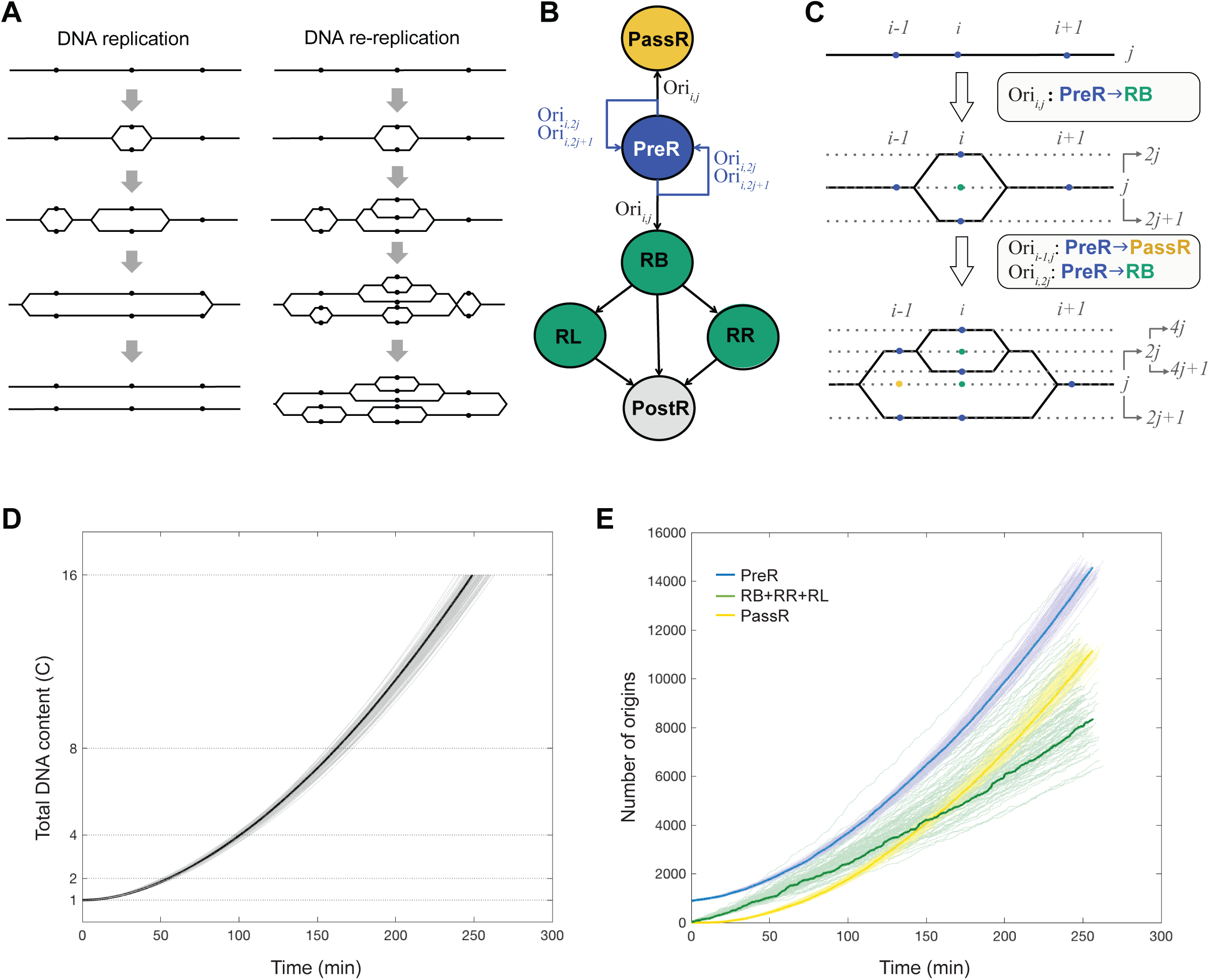
A stochastic hybrid model of DNA re-replication. **A** Normal DNA replication vs. DNA re-replication. Normal DNA replication (left) starts from multiple replication origins (shown here as dots) and is tightly controlled, ensuring that during each cell cycle every origin fires once and precisely two DNA copies are produced. During DNA re-replication (right), re-firing of the origins results in many DNA copies on multiple strands and uneven amplification of the genome. **B** Abstract representation of the DNA re-replication model. Circles of different colors represent discrete origin states and arrows allowed transitions (black: transitions in normal replication, blue: transitions allowed only in re-replication). When an origin fires or is passively replicated, its offspring automatically fall into the PreR state and can thus fire or be passively replicated again. **C** Evolution of re-replication and example transitions between states. Dots of different colors correspond to origins of different states (same as in B). Solid black lines represent synthesized DNA and dotted grey horizontal lines correspond to different strands (strand index shown in grey on the right). Initially, all origins pictured are in the PreR state and located on strand j. Then Ori_i,j_ fires and its offsprings, Ori_i,2j_ and Ori_i,2j+1_ automatically fall into the PreR state. Next, the left fork of Ori_i,j_ reaches the location of Ori_i−1,j_, which leads to its passive replication and the birth of origins Ori_i−1,2j_ and Ori_i−1,2j+1_ that automatically fall into the PreR state. In the meantime, Ori_i,2j_ also fires and creates Ori_i,4j_ and Ori_i,4j+1_ which again fall into the PreR state. Note that the doubling of the strand index (starting with the original strand j = 1) allows us to uniquely identify all strands. **D** DNA re-replication kinetics for 100 Monte Carlo simulations. Total DNA content (C) over time. Different curves correspond to different simulations. **E** Total number of origins per state over time. Different colors correspond to different origin states and different curves to different simulations. Highlighted curves correspond to the simulation closest to the mean.

Contrary to normal DNA replication, where exactly 2 DNA copies are produced, in re-replication origin re-firing allows each origin to produce multiple copies (referred to here as offspring) on multiple resulting strands (Fig. 1A). Each origin copy, whether ancestral or offspring, can be identified by its genomic location and strand index, and is denoted as Ori_*i*,*j*_, where *i* = 1, …, *n* denotes the origin index and *j* = 1, …, *m* the strand index. Fig. 1B pictorially summarizes the discrete dynamics of the model. Similarly to the model of (Lygeros et al., 2008), at any point in time, each origin can be in one of the depicted six states: pre-replicative (*PreR*), replicating in both directions (*RB*), replicating only to the right or to the left (*RR* or *RL*), Passively replicated (*PassR*) and post-replicative (*PostR*). The continuous dynamics are deterministic and model the movement of the replication forks. Uncertainty plays a vital role in re-replication, and it is represented by modeling the time and location of origin firing and re-firing as stochastic events. Transitions between discrete states, depicted as arrows in Fig. 1B, depend on both the continuous and stochastic dynamics of the system. In contrast to normal DNA replication, origins that have already fired or have been passively replicated can re-fire multiple times. These transitions are visualized in an example scenario in Fig. 1C: when an origin fires (transition *PreR* → *RB*) or is passively replicated (transition *PreR* → *PassR*), it generates two origins on two new strands, which automatically fall back into the *PreR* state and can thus fire again (blue arrows in Fig. 1B). To assign firing propensities of these newly-replicated origins, we assume that the total firing propensity (the sum of all firing propensities of every origin in the cell) remains constant (see limiting factor hypothesis (Lygeros et al., 2008) and below for alternative implementations). Each time an origin fires (or is passively replicated), its firing propensity gets dynamically redistributed to all pre-replicative origins across all strands, in proportion to their current firing propensity.

The DNA re-replication model requires the following inputs (Supplemental Fig. S1): (i) total genome length, measured in base pairs, (ii) genomic locations of all origins, measured in base pairs, (iii) intrinsic firing efficiencies of all origins and (iv) fork speed, measured in kilobases replicated per minute (kb/min). Provided that this information is available, the model is applicable to any eukaryotic genome. For the purposes of this work, the model was instantiated for the case of fission yeast (*S. pombe*). This organism has long served as a model system for the study of DNA replication control, as it exhibits conserved features, while its small genome (total genome length ≈ 14 × 10^6^ bases and three chromosomes) simplifies analysis. Exact origin locations and their intrinsic firing efficiencies (fraction of origins that fire when fork movement is blocked with hydroxyurea) have been measured experimentally across the complete fission yeast genome during normal replication (Heichinger et al., 2006). Supplemental Table S1 shows the locations and efficiencies of the 893 fission yeast origins used as input. While initial estimations of fork speed during normal DNA replication were around 3 kb/min (Heichinger et al., 2006; Raghuraman, 2001; Yabuki et al., 2002), recent estimations of fork speed range from 0.5 to 1.5 kb/min (Duzdevich et al., 2015; Sekedat et al., 2010). Mean fork speed in re-replication is expected to be slower than normal replication, due to limited nucleotide pools and interference between forks. Therefore, in our base-case model, we set fork speed equal to 0.5 kb/min and assumed it is uniform across the genome. In contrast to normal DNA replication, where the process is completed when all genomic regions have doubled, in re-replication there is no defined endpoint. The ploidy level *C*, i.e., the total amount of genomic material synthesized with respect to the initial amount, was used to define the stopping time of simulations. It should be noted that re-replication is not expected to progress evenly across the genome and therefore different genomic regions will be amplified to different extents by the end of the simulation.

Monte Carlo simulations of the model for the aforementioned inputs were used to study the re-replication process. Since the model is stochastic, each simulation corresponds to a sample path of the stochastic process, i.e., a distinct sequence of random events; Monte Carlo simulations permit the estimation of statistics over multiple such sequences. Due to the complexity of the process, the discrete and continuous state space quickly becomes very large. In a normal DNA replication cycle, the total number of origins doubles and reaches 1786 as the total DNA content *C* increases from 1 to 2, while the maximum number of active forks will be up to double the number of origins. During the course of re-replication, however, the number of origins and forks, equivalent to the discrete and continuous states of the model, increases drastically: in an example simulation, already by the time that *C* reaches 2, 3861 origins at various states and 2515 active forks are present. For *C* = 16, the count of origins and forks escalates and increases almost 10-fold, with 35259 origins and 18827 active forks present.

In Fig. 1D, examples of the kinetics of DNA synthesis over time are shown. Each curve corresponds to a single simulation, and the spread between individual curves indicates variability due to the stochastic nature of the model. We observe that the increase in DNA content over time is exponential, and a DNA content of 8*C* is reached within approximately 3 hours, in the same range as experimental observations (Kiang et al., 2010; Mickle et al., 2007). The number of active (*RB*, *RR*, *RL*), passive (*PassR*), and pre-replicative origins (*PreR*) over time are shown in Fig. 1E. During the course of re-replication, the number of active and passive origins increases rapidly over time. Initially, passively replicated origins are fewer than the actively replicated ones, but as re-replication progresses, the number of passively replicated origins increases faster and surpasses the number of origins that fired. This indicates that the process gets eventually dominated by passive replication instead of firing events. Pre-replicative origins increase exponentially, as all firing and passive replication events lead to the birth of new *PreR* origins.

We have therefore developed a model which can capture re-replication dynamics across an entire genome, accounting for transitions in origins states, fork movement and stochasticity.

### DNA re-replication at a population level: Model validation

Simulation results from our model were compared to experimental data from re-replicating fission yeast cells. Specifically, we computed *in silico* mean amplification profiles across the genome, referred to as signal ratios in (Kiang et al., 2010), by averaging the number of copies for each origin location and normalizing it to the genome mean in 100 simulations. In these profiles, peaks above 1 correspond to highly re-replicated regions, and valleys below 1 correspond to regions that are under-replicated with respect to the mean. Mean profiles computed at 16*C* are shown in Fig. 2, bottom row. These were compared to re-replication profiles defined experimentally (Kiang et al., 2010), where location-specific amplification was assessed using Comparative Genomic Hybridization (CGH) in fission yeast cells co-overexpressing the licensing factors Cdc18 and Cdt1 (Fig. 2, top row). The profiles appear overall similar, with several peaks coinciding. Indeed, our model predictions fitted experimental observations reasonably well, as the Spearman correlation coefficient *ρ* between experimental and simulated whole-genome re-replication profiles was statistically significant for all three fission yeast chromosomes (ρ=0.6 and p-value=3.6*10_-41_ for Chromosome I, ρ=0.61 and p-value=5.7*10_-33_ for Chromosome II, and ρ=0.5 and p-value=7.3*10_-12_ for Chromosome III). To better compare simulated and experimental profiles, a peak-calling algorithm was used, which identified 29 and 22 peak locations in experimental and simulated profiles respectively (dotted vertical lines), representing regions of amplification in a population of re-replicating cells. Details on the denoising and peak finding processes are given in Methods, and indices and locations of both peak sets are given in Supplemental Table S2.

**Figure 2.**
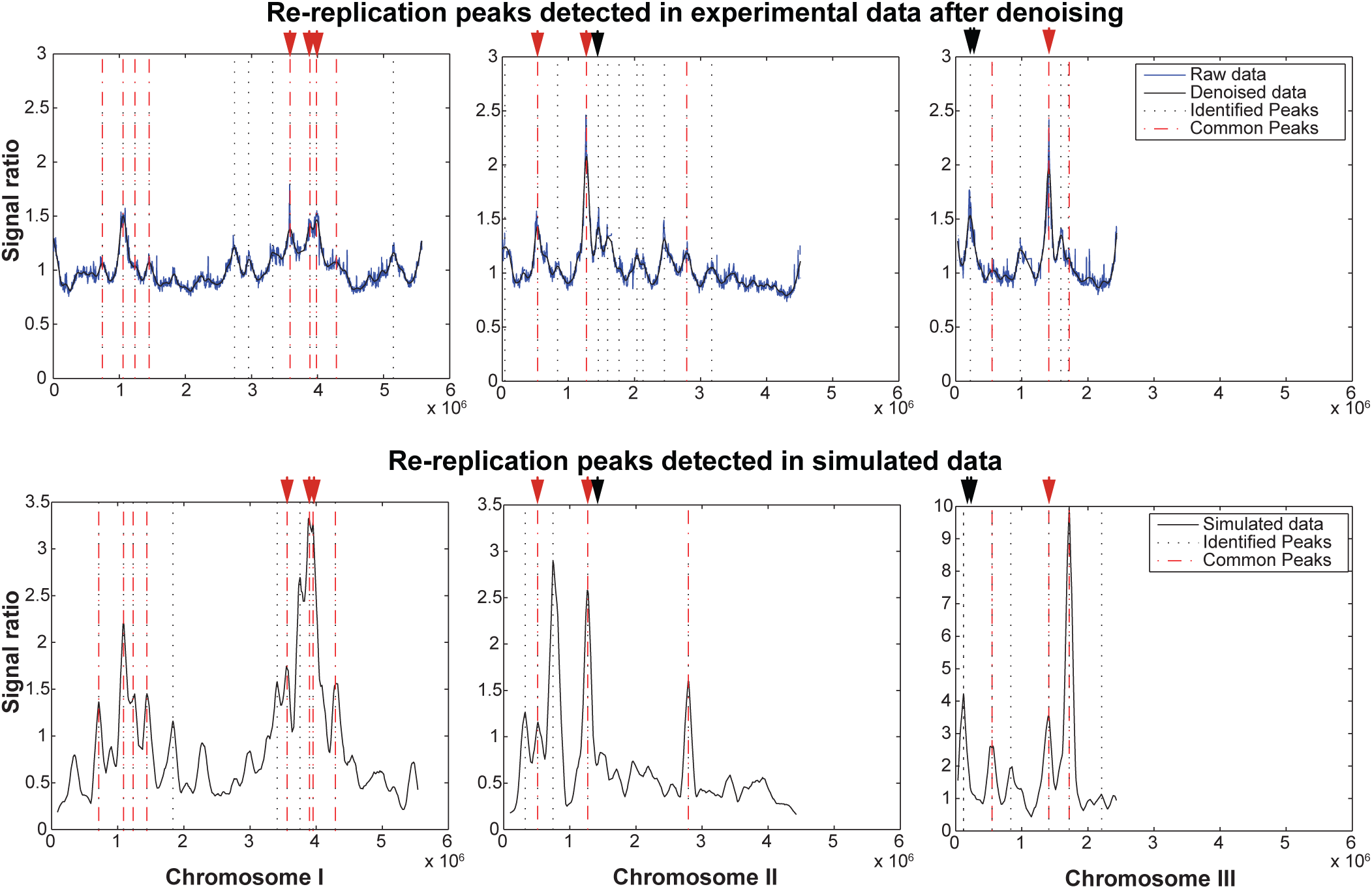
Analysis of in silico data at a population level. Comparison between experimental (top row) and simulated (bottom row) mean amplification profiles for 100 Monte-Carlo runs for 16*C* along the three chromosomes of the fission yeast genome. Identified peaks, representing re-replication hotspots, are marked in dotted vertical lines. Common peaks between simulated and experimental data are marked in red dashed vertical lines. Peaks corresponding to the 9 amplification regions identified by (Kiang et al., 2010) are marked in vertical arrows (red: identified also in simulated data, black: identified only in experimental data).

We observe that most peaks predicted from simulations correspond to major amplification peaks in the experimental data: out of the 22 re-replication peaks in the simulated data, 14 also exist in the experimental data (precision ≈ 0.64). To further assess this agreement between experimental and simulated peaks and examine whether it could be attributed to chance or our peak-matching algorithm parameters, we repeated the analysis 100,000 times using a null model of 22 randomly picked genome locations (Methods). We found that the median overlap between experimental and random peaks was 2 and the maximal overlap, occurring only once in all 100,000 random repetitions, was 10, indicating that the probability of the model’s prediction being attributed to chance is less than 1 in 100,000. Moreover, out of the 9 amplification regions identified in (Kiang et al., 2010), 6 are also present in the simulated profiles. Some inconsistencies do exist between the two datasets: out of 29 peaks in the experimental data, 15 are not predicted (false negative rate ≈ 0.52). Most of these false negatives however are attributed to peaks that are present but appear less sharp in the simulations (e.g., peaks in the middle part of Chromosome II) or to minor disagreements due to linear shifts in peak locations (e.g., the first and third true peaks of Chromosome III). These minor inconsistences could potentially be attributed to the origin efficiencies used as input (Heichinger et al., 2006). Indeed, comparison of the input data with a different set of origin location and efficiencies, estimated using a polymerase usage sequencing (Pu-seq) strategy during an unperturbed cell cycle in *S.pombe* (Daigaku et al., 2015), revealed minor disagreements across datasets, which could account for some of the observed inconsistencies (data not shown). Striking inconsistencies are relatively few, notably the 5th peak of the simulated data in Chromosome III that has a large difference in intensity and the subtelomeres that are highly amplified in experimental data but not in the simulations. These point to location-specific effects, not explicitly specified in our model, as previously suggested for subtelomeric regions (Mickle et al., 2007). Such isolated events however do not significantly affect the overall re-replication dynamics.

We conclude that simulated population data fit experimental data genome-wide reasonably well, validating our approach.

### Sensitivity analysis

We next sought to investigate the effects of different model parameters and assumptions. We first varied fork speed: 0.5 kb/min (base case) was compared to 1 and 3 kb/min (Supplemental Fig. S2). As expected, the increase in DNA content progresses faster at higher fork speeds: *C* = 8 is reached at 164, 121 and 74 min as fork speed increases from 0.5 to 1 and 3 kb/min (Supplemental Fig. S2A, B), with estimates for 0.5 kb/min closer to experimental observations. Passive replication becomes more dominant as fork speed increases (Supplemental Fig. S2C).

A second variant of the model was tested, that differs on how the firing propensities of newly-replicated origins are assigned. In this variant (referred to as Unlimited Factor or UF) we assume that when an origin fires or is passively replicated, the offspring inherit the same firing propensity as the parent. This implies that firing propensities depend only on the genomic location and hence remain the same for the same origin across all strands. Under this assumption the total firing propensity (i.e., the sum of the propensities of all the origins) will increase during re-replication (Supplemental Fig. S2D). By contrast, in the base-case (referred to hereafter as Limiting Factor or LF), the total system firing propensity remains constant during re-replication while the firing propensities of individual origins decrease, as more origins are born (Supplemental Fig. S2E). The two variants of the model reflect different biological hypotheses. The UF variant represents a situation where all factors needed to license and activate an origin are available in virtually unlimited quantities. The LF variant represents a situation where one or more of these factors exists in limited quantities and binds to origins proportionally to their intrinsic efficiencies (Lygeros et al., 2008; Rhind, 2006).

Simulation kinetics for the UF variant at a fork speed of 3 kb/min are shown in Supplemental Fig. S2E. When unlimited copies of an activation factor (UF variant) are assumed, the process is fast, as DNA content doubles approximately every 12 minutes and reaches 16*C* in less than 1 hour. For the LF variant, on the other hand, each doubling needs gradually more time to complete, and re-replication requires roughly twice as much time to reach the same *C* levels as in the UF case. In the UF model, the number of passive and active origins increases rapidly with a comparable count (Supplemental Fig. S2C), in contrast to the LF model where active origins increase at a slower rate and are eventually outnumbered by the passive ones. The re-replication process predicted by the UF model is much faster than experimentally observed (Supplemental Fig. S2B), suggesting that unlimited re-replication is unlikely to take place within cells.

Mean amplification profiles across the genome for different values of fork speed are shown in Supplemental Fig. S3 at 16*C*. Profiles appear flatter as fork speed increases, consistent with increased passive replication. The sites of over-amplification however appear at similar locations along the genome, suggesting that re-replication dynamics along the genome are robust to varying fork speeds. In Supplemental Fig. S3, amplification profiles genome-wide are also compared between the base-case model (LF variant, fork speed of 0.5 kb/min) and the UF variant (fork speed of 3 kb/min). We observe that both profiles follow a very similar pattern, with the UF profiles characterized by somewhat sharper peaks, indicating more firing from the underlying origins. Importantly, all amplification peaks shown in Supplemental Fig. S3 are consistently present in both model variants and parameter values.

We conclude that the base-case model is robust to model assumptions.

### Single-cell analysis: heterogeneity across the genome

Re-replication across the genome has only been studied so far at the population level. Although population-based methods enable the exploration of global characteristics, they mask the underlying variability at a single-cell level, as only the most prominent regions “survive” in the mean amplification profiles. The model described here permits analysis of cell-to-cell heterogeneity of the amplification levels across the genome, as each simulation corresponds to a distinct sequence of events taking place within one cell.

To assess variability at the single-cell level, we compared amplification plots from single simulations, generated by the base-case model at 2C and 16C. Single-cell profiles are characterized by a high degree of variability and can deviate significantly from the mean behavior (examples of four random simulations at 16C in Supplemental Fig. S4). To quantify the variability in the simulations genome-wide, we used the Shannon entropy, an information-theoretic metric (Methods) on discretized data from 100 simulations, where 0 and 1 correspond to copy number levels less and more than the genome mean, respectively. As shown in Supplemental Fig. S5, this analysis indicates that whether an origin is amplified or not is highly unpredictable at 2*C*, while at 16*C* the entropy becomes bimodal, with half of the origins consistently over- or under-replicated.

In Fig. 3, a zoom in on 1Mb of Chromosome I is shown at 2*C* (Fig. 3A) and at 16*C* (Fig. 3B) for the same four randomly selected individual simulations as in Supplemental Fig. S4. We observe that amplification levels along the genome vary across the simulations, pointing to a high degree of heterogeneity. This variability is especially prominent early on in the process (2*C*), where certain origins have been amplified to a high degree, while most of the genome remains normal. We analyzed the copy number levels at 16*C* of an individual origin in this region (red dot in Fig. 3A-B), for which a high degree of heterogeneity was not apparent in the particular simulations selected. We observed that, when looking at all simulations, the distribution (Fig. 3C) is right-skewed with a heavy tail, showing that high variability in copy number levels is indeed present.

**Figure 3.**
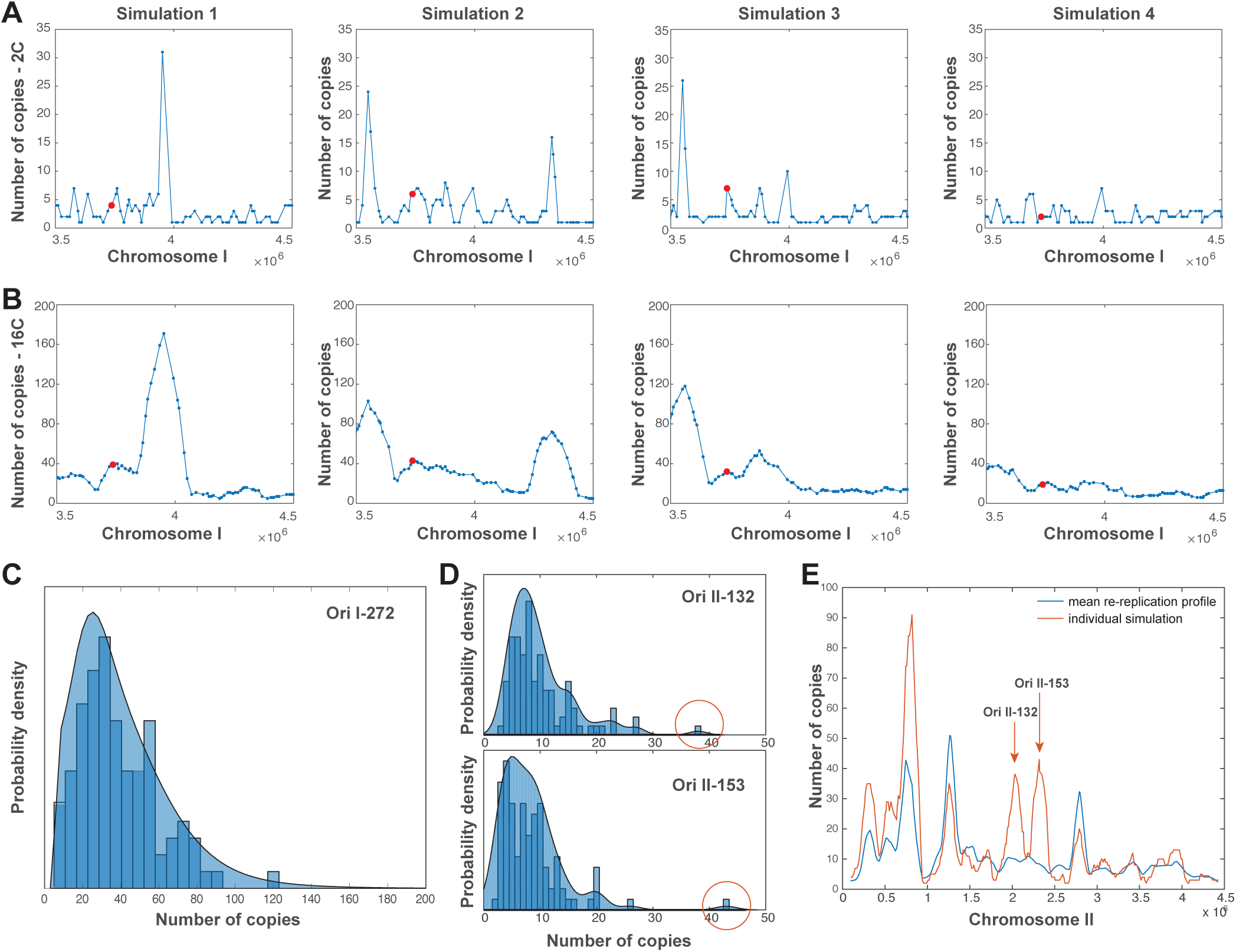
Analysis of in silico data at a single-cell level exposes heterogeneity. **A-B** Model simulations expose heterogeneous patterns of re-replication at a single-cell level. Individual simulations of the stochastic model lead to markedly different amplification levels. Shown here are number of copies for all origins (marked in circles) on a random region of Chromosome I, resulting from four random simulations of the model at a total DNA level of 2*C* (A) and 16*C* (B). **C** Copy number distribution of one individual origin of Chromosome I (Ori I-272), highlighted in red in (A-B), from the 100 simulations at 16*C*. **D** Distributions of copy number levels of weak origins Ori II-132 and Ori II-153. Outliers of the distributions are marked in circles. **E** Mean amplification profile of Chromosome II from all 100 simulations (blue) vs. single-cell amplification profile corresponding to one individual simulation for which Ori II-132 and Ori II-153 are amplified.

To determine whether every region along the genome is amenable to amplification upon re-replication, we computed the number of times each origin was amplified more than the genome mean (*C*=16). This analysis showed that 739 out of the total 893 origins are amplified above the mean at least once in the set of 100 simulations analyzed. This suggests that even regions of low efficiency can potentially be amplified. A striking example is given in Fig. 3D-E for origins Ori II-132 and Ori II-153. As seen from their copy number distributions (Fig. 3D) and the mean re-replication profile (Fig. 3E), both origins are under-represented in the population and reside in a region of almost no re-replication. However, as seen in the single-cell profile, they can potentially re-replicate, and yield copies high above the population mean. This suggests that under re-replication, multiple combinations of co-amplified regions will appear, even for low efficiency regions.

We conclude that re-replication can drive different genomic regions to be amplified in different cells, leading to heterogeneity at the single cell level.

### Single-cell heterogeneity observed *in vivo*

To experimentally explore the cell-to-cell copy number variability under re-replication *in vivo*, the relative amplification level at a specific genomic region was assessed using the LacO/LacI system. Specifically, high affinity binding of an ectopically expressed, fluorescent-tagged lactose inhibitor (lacI-GFP) onto stably integrated lac operator (lacO) arrays allows the visualization of a targeted genomic region as a fluorescent dot, the intensity of which reflects the copy number of the lacO-targeted region (Kitamura et al., 2006). To induce re-replication in a controllable manner, a fission yeast cell strain bearing a truncated form of the licensing factor Cdc18 (*d55P6-cdc18*) under the repressible promoter nmt1 was employed (Baum, 1998). Absence of the vitamin B1, thiamine, activates the promoter and leads to Cdc18 overexpression and medium re-replication levels, confirmed by FACS analysis in Supplemental Fig. S6A. Additionally, the same strain carries the *lys1+* locus marked by the lacO-lacI system (Fig. 4A) (Petrova et al., 2013). The *lys1+* gene is located between Ori I-272 and Ori Ι-273, which present 60% and 39% efficiency, respectively. Copy number levels for Ori I-272 in individual simulations and across the whole population are shown in Fig. 3A-C.

**Figure 4.**
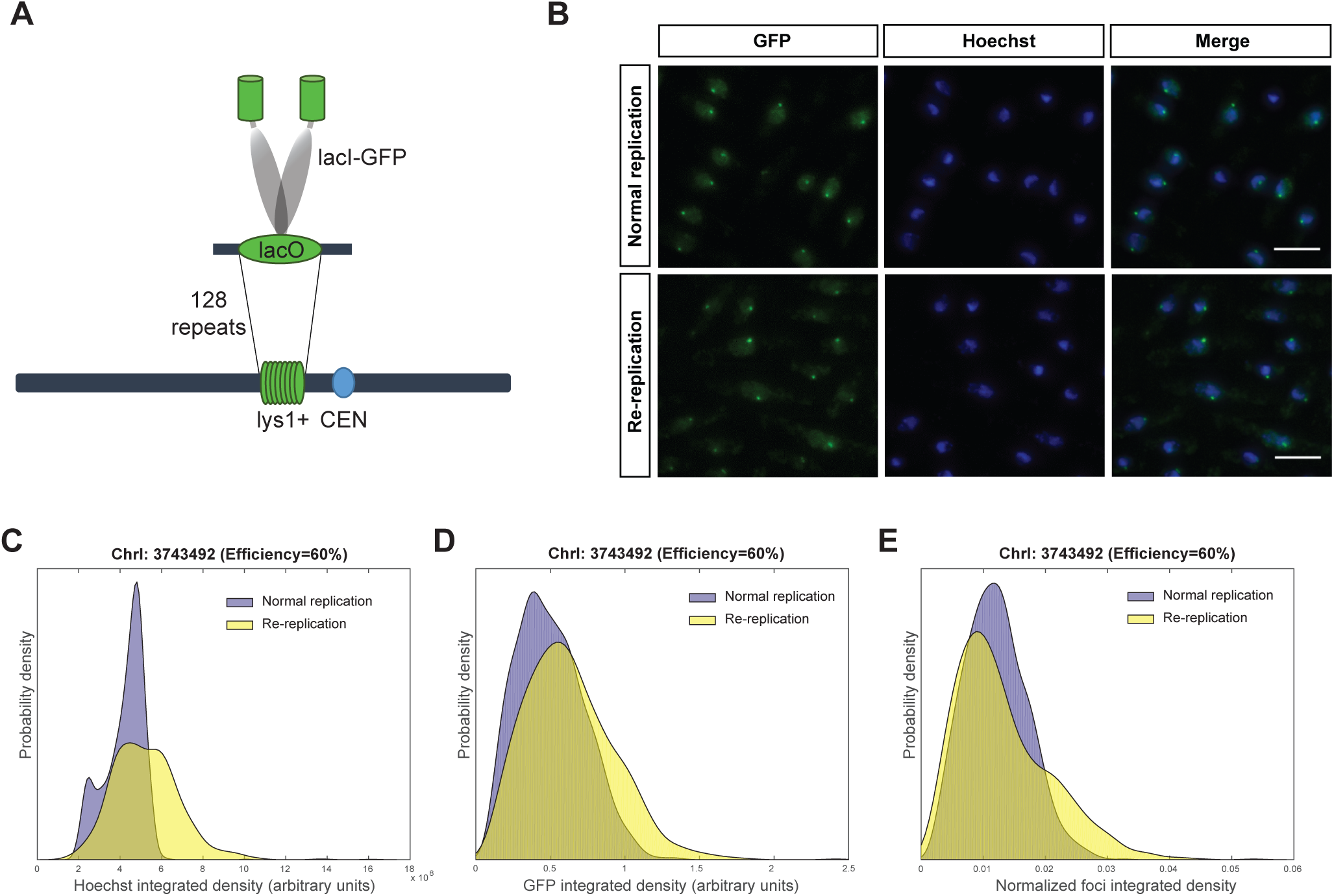
Quantification of lys1+ region in normal and re-replicating conditions. **A** A region proximal to lys1+ gene (ChrI: 373492) was labelled by integration of 128 LacO repeats in fission yeast cells expressing LacI-GFP. **B** Visualization of GFP labelled region in cells growing under normal replication and re-replication. Scale bar: 5 μm. **C-E** Histograms of Hoechst, GFP integrated density and normalized foci integrated density in cells undergoing normal replication (n=1632) and re-replication (n=1234) are presented.

Re-replication was induced for 30 hours at 25 ºC, or not as a control, and the cells were fixed, stained with the DNA dye Hoechst and imaged in a widefield epifluorescence microscope (Fig. 4B). Image analysis revealed an increase in DNA nuclear staining in the re-replicating cells (Fig. 4C), as well as, an increase in the intensity of the lacI-GFP dot (Fig. 4D), each of them indicating increased genomic content and increased copies of the *lys1+* locus under re-replication, respectively. As shown in Supplemental Fig. S6B, in control cells the distribution of DNA nuclear staining is consistent with the presence of G1, S and G2 phase cells, with G2 cells having approximately double the DNA content of G1 cells, while lacI-GFP foci intensities correlate with the DNA content. On the contrary, re-replicating cells does not present clear separation of populations and varying levels of re-replication are observed in different cells. Foci intensities appear to vary independently of the DNA content. To estimate the relative copy number of the *lys1+* region with respect to the DNA content at the single cell level, the intensity of each GFP dot was normalized with the total DNA nuclear intensity in each individual cell (Fig. 4E). We observe that under re-replication the distribution of the normalized GFP intensity is positively skewed with a long tail and an increased coefficient of variation compared to the normal replicating sample (41.18% for normal and 57.54% for re-replication), in agreement with the simulated data at this region (Fig. 3C). We conclude that cell-to-cell heterogeneity in the number of copies of the *lys1+* genomic locus is evident in fission yeast cells undergoing re-replication, consistent with *in silico* analysis.

### Rules governing DNA re-replication across the genome

#### Intrinsic origin properties

To unveil the rules which dictate which regions will become amplified along the genome, we first assessed the dependence of amplification levels of individual origins on their intrinsic efficiencies, as determined experimentally (Heichinger et al., 2006). Fig. 5A shows histograms of the copy number distributions from simulations of the LF model at 0.5 kb/min at 16*C* of two origins (Ori II-45 and Ori II-54), with high and low efficiencies (62% and 9% respectively (Heichinger et al., 2006)). The median number of copies of each origin is consistent with its efficiency (notice again the positively skewed distribution with long tails discussed above). The scatterplot of Fig. 5B shows a strong correlation between mean number of fires at 16*C* and efficiency for all origins (Spearman correlation coefficient - *ρ* = 0.96). The coefficient of variation (ratio of standard deviation over the mean) is inversely correlated to the efficiency (*ρ* = −0.89), with weak origins showing much higher variation than strong ones. Mean number of copies of each origin show a weaker correlation to firing efficiency (Fig. 5C, *ρ* = 0.4) and a coefficient of variation weakly linked with efficiency (*ρ* = 0.12). Interestingly, a higher spread at low efficiencies is observed, with origins considered dormant (efficiencies less than 10%) occasionally significantly amplified with respect to the population mean. Last, a scatterplot of the mean number of passive replications vs. the efficiency (Fig. 5D) indicates a much weaker correlation (*ρ* = 0.17) and a weak efficiency-related variability (*ρ* = 0.13). This analysis shows that re-firing of a given origin is strongly affected by its efficiency, while additional properties govern levels of amplification of individual loci, which are especially prominent for low-efficiency origins.

**Figure 5.**
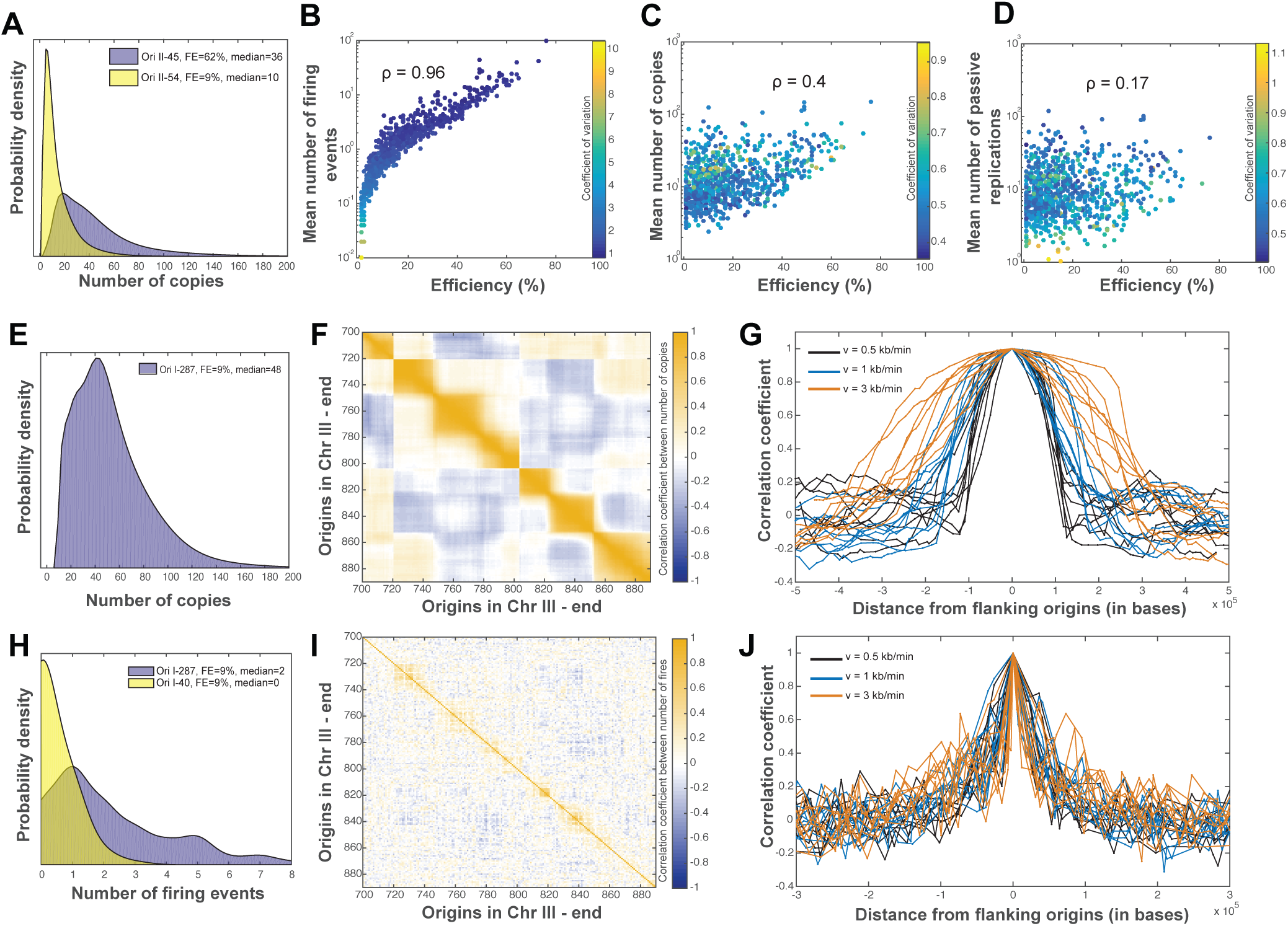
Analysis of in silico data at a single-cell level linked with intrinsic properties and points to in cis effects. Simulation results from 100 Monte Carlo simulations of the LF model at 0.5 kb/min at 16*C*. **A-D** Amplification levels of individual loci with respect to their intrinsic properties. **A** Distributions of copy number levels for two origins of high (purple) and low (yellow) efficiency (origin indices, efficiencies and median number of copies given in the legend). **B** Scatterplot of mean number of fires vs. firing efficiency for all origins shows a strong correlation between firing events and firing efficiency (Spearman correlation coefficient value - ρ = 0.96). Color indicates coefficient of variation (standard deviation/mean), and points to higher variability in firing for the weak origins. **C** Scatterplot of number of copies of individual origins vs. their firing efficiencies shows a weaker correlation (ρ = 0.4) and variation less dependent of efficiency. **D** Scatterplot of mean number of passive replications vs. firing efficiency shows a very low correlation (ρ = 0.17). **E-G** Amplification levels of individual loci with respect to local effects. **E** Distribution of copy number levels of weak origin I-287, residing next to strong origin I-288, shows elevated levels due to passive replication. **F** Heatmap of correlations of copy number levels between different origins exposes strong in cis effects, as shown here for a zoomed in region in the end of Chromosome III. Color indicates Spearman correlation coefficients. **G** Correlation coefficients between copy number levels of the 10 origins with the highest intensity (different lines) and their neighboring origins, centered and zoomed in the origin locations. Different color indicates varying values of the fork speed. **(H-J)** Firing activity of individual origins with respect to local effects**. H** Distributions of firing events for two origins of low efficiency, with efficient (purple) and inefficient (yellow) neighbors, shows that weak origins fire more often when residing next to strong ones. **I** Same as in (F) but showing correlations between number of fires across the genome. **J** Same as in (G) but showing correlations between number of fires of prominent origins and their neighbors.

#### In cis effects

Next, we investigated origins whose amplification levels could not be explained merely by their intrinsic efficiency. A relevant example is given in Fig. 5E; from the distribution it is clear that, although Ori I-287 has a very low efficiency, its amplification levels are much higher than expected. A closer examination of the neighboring origins reveals that its left-flanking origin (Ori I-288) is one of the most efficient in the genome, with a firing efficiency of 73%; their copy number levels are strongly correlated (*ρ* = 0.98). To further investigate this, we computed correlation coefficients across copy number levels of all origins in the genome (whole genome in Supplemental Fig. S7 A-B, zoom in Chromosome III – end in Fig. 5F). This analysis indicated strong correlations between adjacent origins, pointing to *in cis* effects. To better understand the extent of this effect, we computed correlations between copy number levels of the 10 peaks with the highest amplification and their neighborhood (Fig. 5G). Since our previous analysis showed that fork speed affects the extent of passive re-replication, we also computed correlations using the simulations with a fork speed of 1 kb/min and 3 kb/min. The results reveal that copy numbers of each central amplification origin are significantly positively correlated with the ones of its right and left flanking up to a distance of 0.1 megabases; it is also clear that as speed increases, the extent of positive correlation increases as well and for fork speed equal to 3 kb/min it reaches a distance of 0.5 megabases.

We then asked how the firing activity of individual origins is affected by the activity of its neighbors. We focused our analysis on Ori I-287 and Ori I-40, two origins that share the same low efficiency (9%), but Ori I-287 has an immediate neighbor with high efficiency (73%) whereas Ori I-40 does not. We observed that Ori I-40 does not fire the majority of times, whereas Ori I-287 appears more active and occasionally fires even more than 5 times (Fig. 5H). We then followed the same methodology as above and computed the correlation coefficient between the number of fires of different origins across the genome (Fig. 5I). This analysis indicated that indeed local effects exist, suggesting that the more times an origin fires, the more will its neighbors fire as well. We notice that the firing events of each central amplification origin are significantly positively correlated with the ones of its immediate right and left flanking neighbors, however, this time the correlation spans a smaller region, drops sharply with distance from the central origin and does not appear affected by fork speed (Fig. 5J). These findings indicate that, in addition to passive re-replication, *in cis* effects between adjacent origin locations are also implicitly attributed to increased firing activity of weak origins located close to strong origins. Early firing of a strong origin will increase the newly born copies of a nearby weak origin, facilitating its re-firing.

#### In trans effects

To explore the variability of the re-replication process genome-wide, we performed a principal component analysis of the genome-wide amplification profiles of 100 simulations at 16*C* and visualized the results as a biplot of the first two principal components (Fig. 6A), where dots correspond to simulations and vectors indicate the PCA loadings, i.e., the correlation of each origin to the unit-scaled first two principal components. From this it becomes clear that a large amount of the variability in the simulations is dominated by two different origins of Chromosome III (Ori III-11 and Ori III-118). Specifically, the first and second principal component correlate strongly with Ori III-118 and Ori III-11 respectively, while Ori III-11 additionally appears to correlate negatively with principal component 1.

**Figure 6.**
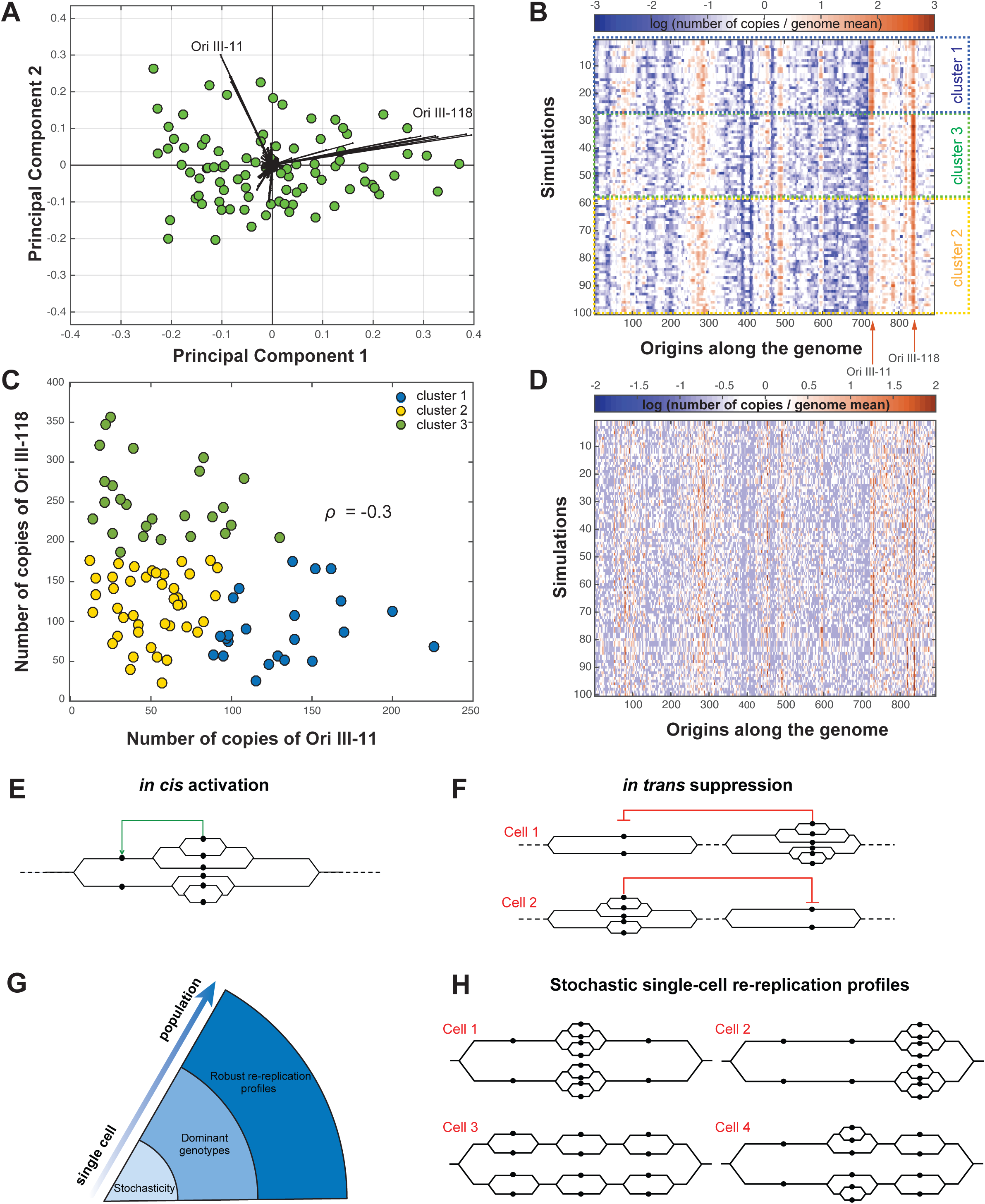
Analysis of in silico data at a whole genome level points to in trans effects within the genome. **A** Variability of copy number levels genome-wide is governed by prominent origins. Results of a PCA analysis of the in silico copy number data, shown as a biplot of the first two principal components. Dots correspond to simulations and black vectors expose each origin’s contribution to the first two components, both in terms of magnitude and direction (marked here for the two most prominent ones). **B** Heatmap of DNA content (rows: simulations, columns: origins) for 100 simulations at 16*C* after clustering with a k-means algorithm and k=3. Color indicates DNA amplification levels, expressed as the log ratio of individual vs. genome mean number of copies. Identified clusters are marked with different colors. **C** Scatterplot of number of copies for origins Ori III-11 and Ori III-118 shows a negative correlation (ρ = −0.4). Colors correspond to simulations belonging to each of the three clusters identified in B. **D** Evolution of re-replication over time. Heatmap of DNA content for simulations of (B) at an earlier DNA content of 2*C* shows no cluster-specific patterns at a low-re-replication context. **E** Underlying characteristics of DNA re-replication. In cis effects between adjacent loci. Passive re-replication of inactive origins from their efficient neighbors leads to increased copy numbers and implicitly increases their firing activity. **F** In trans effects between distant loci. Increased amplification of one locus leads to in trans suppression of a distant locus. **G** Emerging properties of DNA re-replication, depending on the level of analysis. **H** In silico re-replication profiles. Simulation results reveal many possible genotypes within a population, shown here in a schematic view for three hypothetical origins. Although the total DNA content is the same in all four single cells, individual copy number levels vary greatly.

To further explore how specific origins may affect genome-wide amplification profiles, we clustered profiles using *k*-means clustering, estimated the optimal *k* using the Gap statistic (Tibshirani et al., 2001) and the stability of the clustering using the Adjusted Rand Index (ARI) (Hubert and Arabie, 1985) (details in Methods). For 100 simulations at 16*C* an optimal number of *k* = 3 clusters was identified, and the cluster assignments were very consistent across 100 random initializations, with a mean ARI of 0.95 (standard deviation = 0.05). These clusters correspond to 3 groups of simulations characterized by different patterns of re-replication at a genome level (Fig. 6B). The clusters appear to be dominated by the amplification of origins Ori III-11 and Ori III-118 in a mutually exclusive manner: either one of the two origins is amplified (clusters 1 and 3) or they are both relatively low (cluster 2). Indeed, as shown in Fig. 6C, levels of amplification of Ori III-11 and Ori III-118 are negatively correlated in individual simulations (*ρ* = −0.3, p-value=0.0025) and characterize the 3 clusters. Taken together, these findings indicate *in trans* effects within the genome.

We next examined the same simulations at a DNA content of 2*C* (Fig. *6D*, same ordering as in Fig. 6B). We observe that while specific amplification regions are starting to emerge, the re-replication levels for the majority of the genome are around the genome mean, amplification occurs in random regions along the genome and the process is governed by a high degree of variability. At the same time the difference between single-cell profiles of the previously identified clusters is not noticeable. To validate this, we went on to independently cluster the data and estimated an optimal *k* of only 1 cluster. Forcing *k* equal to 3 and estimating the stability across 100 random initializations indicated close to random cluster assignment between different runs (ARI = 0.36 ± 0.15). Comparing the clusters found for 2*C* and 16*C* also indicated very low agreement (ARI = 0.34 ± 0.10). Last, the correlation coefficient between Ori III-11 and Ori III-118 for 2C is now non-significant (*ρ* = −0.09, p-value=0.36).

We conclude that re-replication is initially characterized by a high degree of randomness, while *in trans* effects become evident as the re-replication process progresses, leading to preferred genome-wide patterns of re-replication at high DNA content. These are dominated by a small number of high activity origins, whose amplification to high levels is mutually exclusive.

## Discussion

### A stochastic hybrid model of DNA re-replication

In this work a stochastic hybrid model of DNA re-replication was presented, developed by refining existing work of normal DNA replication so that it allows for origin re-firing. The model accurately portrays the interplay between discrete dynamics, associated with different origin states, continuous dynamics, associated with the movement of the replication forks, and stochasticity, associated with random firing and re-firing events. Transitions between discrete states depend on both the continuous and stochastic dynamics of the system, such as firing events or merging of neighboring forks. In addition, two automatic transitions, specific to the re-replication case, are incorporated in the model: when an origin fires or is passively replicated its descendants automatically fall into the pre-replicative state and can potentially fire or be passively replicated again.

Using input data from experimentally determined origin locations and efficiencies from fission yeast, the model allows the simulation of re-replication along the complete fission yeast genome and thus the exploration of re-replication kinetics genome-wide. Two alternative variations of the model have been implemented, depending on how the firing propensities of the newly-born origins are assigned. In the base-case model variation (LF model), the total system propensity is kept constant and continuously redistributed to all existing and newly born origins. In the alternative variation (UF model), offspring inherit the same firing propensity as the parent, and as the total number of origins increases exponentially, the total system firing propensity will also increase. Experimental evidence in support of such a limiting factor hypothesis points to initiation factors that participate in origin activation (Aparicio, 2013; Mantiero et al., 2011; Patel et al., 2008; Wu and Nurse, 2009). In (Lygeros et al., 2008) it was shown that, for normal replication, redistribution of a limiting factor increases the efficiency of remaining origins and may help explain away the so-called random gap problem (Hyrien et al., 2003).

### Parameters affecting re-replication dynamics

Comparison of *in silico* data for both model variations and experimentation with different values of the model inputs has permitted insight into the model parameters affecting re-replication dynamics. Our analysis indicated that the simulated re-replication completion times were consistent with experimental observations when using the model variation with limiting factor and a fork speed of 0.5 kb/min. This indicates that, as expected, fork speed is slower in re-replication that in normal mitosis, where experimental estimates in yeast vary between 1.6 and 3 kb/min. Since re-replication is a non-physiological process, differences in fork speed could be attributed to various mechanisms, such as activation of checkpoint proteins that stall the forks, impediments caused by fork collisions (Alexander and Orr-Weaver, 2016) and limitations in the amounts of various necessary substrates like dNTPs (DNA building blocks), activation factors etc.

Further exploration using different model variants indicated that when no limiting factor is assumed, the rate of increase in DNA content far exceeds experimental observations. At the same time, in the model variation with limiting factor, re-replication dynamics are dominated by passive replication instead of firing events, whereas in the variation without limiting factor, firing and passive replication contribute equally to the increase in DNA content. Sensitivity analysis using different values of fork speed showed that, when fork speed is decreased, more time is needed to reach the desired DNA content. At the same time an apparent trade-off between fork speed and firing events was noticed, since faster forks resulted in less firing and allowed passive replication to dominate the increase in DNA content.

### Genome-wide profile of re-replication

Analyzing the simulated data at a population level, it is clear that re-replication is non-homogeneous along the genome, as specific regions are preferably amplified and appear as emerging peaks above the genome mean, whereas others appear dormant and under-represented, an observation that is in accordance with existing experimental findings (Kiang et al., 2010; Mickle et al., 2007). Although the model is stochastic, the most highly amplified regions appear to be highly robust with respect to different model variations or different values of the fork speed. By comparing the simulated vs. the experimental amplification profiles, we observe that overall the simulated data reproduce the experimental re-replication pattern on a whole-genome scale, validating our approach. The best-fitting parameter set proved to be when using the model variation that assumes the existence of limiting factor and a fork speed of 0.5 kb/min, reconfirming the previous findings.

Our analysis showed that most highly amplified regions in experimental data are predicted when using the simulations, with some inconsistencies attributed to minor linear shifts or differences in intensity. Striking differences regard specific regions, with the most prominent ones being the subtelomeres, regions highly amplified in experimental data. This difference could be attributed to location-specific mechanisms, such as the suppression of the telomeric origins in normal DNA replication by telomere-associated proteins Rif1 and Taz1. Distorted nuclear architecture during re-replication or limiting abundance of Rif1/Taz1 could lead to the subtelomeric origins escaping their normal control mechanism and getting amplified above the levels that are expected from their experimentally determined mitotic efficiencies (Heichinger et al., 2006).

### Factors that affect amplification levels of individual loci

Analysis of simulated data at a single-cell level permits a more detailed insight into DNA re-replication. Amplification levels of individual loci are primarily affected by intrinsic properties, since copy numbers were found to be highly correlated with firing efficiencies. At the same time, we found that individual origins are able to act *in cis* and amplify the copy numbers of their neighbors. Positive correlation between amplification levels of adjacent loci is primarily attributed to passive re-replication, as forks emanating from the firing origins to both directions passively replicate the left and right flanking origins. At the same time, as the forks create new copies of the passively replicated origins, these newly-born copies can potentially fire again, thus increasing the overall probability of firing events from the neighboring origins. This means that *in cis* elements contribute to amplified copy numbers not only directly by passive re-replication, but also implicitly through increasing the firing activity of their neighbors (Fig. 6E). This type of positive correlation between adjoining regions is a key characteristic of re-replication and serves as a mechanism for indirect amplification of individual loci.

At the same time genome-wide analysis of the single-cell re-replication profiles revealed groups of similarly amplified simulations, characterized by different patterns of re-replication. These patterns were characterized by amplification of specific regions, residing in non-proximal locations across the genome that appeared to exist in opposition. The amplification levels of these loci were found to be negatively correlated and allowed for a clear separation of the clusters. These findings point to *in trans* interactions between distant regions within the genome, suggesting a mechanism for the suppression of the amplification levels of individual loci (Fig. 6F). Such *in trans* negative regulation of distant origins could be explained by competition for the same limiting factor: high-level amplification of a given locus recruits high levels of the limiting factor, indirectly inhibiting firing of other genomic regions.

### Emerging properties of re-replication, revealed by different levels of analysis

Depending on the level of analysis (from the single-cell towards the population level), different properties of re-replication are revealed. At the single-cell level a large degree of heterogeneity is observed, not only in the variations of individual loci among the population, but also in the variability of single-cell profiles. When clustered, different amplification patterns are recognized in these profiles, corresponding to dominant genotypes within the population. Last, at a population level, re-replication appears highly robust and amplification hotspots appear independent of changes in parameters (Fig. 6G). In conclusion, heterogeneity and robustness appear as key players in the re-replicating process that co-exist and act in parallel. This implies that experimental observations of re-replication, based on population-level data, possibly mask the underlying variability in the behavior of single-cells in a population.

### Cell-to-cell heterogeneity leads to genome plasticity

Stochasticity lies at the heart of re-replication: it gives rise to heterogeneous single-cell profiles that correspond to diverse genotypes within the population. Although amplification levels of individual loci within the population are affected by intrinsic properties, *cis* and *trans* acting elements as mentioned above, great variations in copy numbers of individual loci within a population are revealed, evident by skewed distributions with long tails. As each simulation of the stochastic model corresponds to a single cell in a population, each simulated re-replication profile portrays a unique sequence of firing and re-firing events and corresponds to different genotypes within the population (Fig. 6H). By exploiting this property of the model, our analysis showed that, although re-replication profiles at the population level are robust, at the single-cell level they are heterogeneous and can deviate significantly from the mean. Early firing of an origin in a given cell will increase the probability of a second firing event in the same locus (as more strands are born), leading to a positive feed-back loop which will amplify different loci in different cells.

At the same time, using a population size of 100 simulations, the majority of the genome was found to be amplified above the genome mean at least once. This suggests that even regions of low efficiency can potentially be amplified, and that re-replication can, with varying probability, occur anywhere in the genome and generate many diverse genotypes within a population. If we consider that a small colony of yeast cells contains millions of individual cells, it becomes apparent that re-replication can lead to the appearance of amplification events in a variety of chromosomal regions or combinations of regions. These observations indicate that cell-to-cell variability is inherent in re-replication and can lead to a high degree of genome plasticity. By tracking the evolution of single-cell profiles from a low to a high re-replication context, we found that cell-to-cell variability is more prominent at the onset of re-replication, when single-cell profiles appear highly stochastic in nature. As DNA content increases, cell-to-cell variability is less apparent and specific amplification regions dominate the process. At the same time, distinct patterns gradually emerge, representing dominant genotypes within the population that appear to act antagonistically.

### Genome plasticity and possible implications for oncogenesis

Variations in the number of copies of specific genomic loci have long been implicated in the initiation and progression of cancer. For example, oncogenes and genes conferring resistance to drugs have been shown to be frequently amplified in various cancers (Hills and Diffley, 2014; Masood and Bui, 2002; Ross et al., 2004; Santarius et al., 2010; Schmitt et al., 2016). Re-replication, and the resulting increase in the copies of specific genomic loci, could be a mechanism leading to gene amplification (Black et al., 2013; Green et al., 2010). Studies using cancer genome data correlate replication timing with mutation rates during cancer and suggest that early replication is correlated with gene amplifications whereas late replication with copy number losses (De and Michor, 2011; Miotto et al., 2016; Sima and Gilbert, 2014). An overwhelming amount of experimental evidence supports a high level of heterogeneity in cancer cell populations (Burrell and Swanton, 2014; Greaves and Maley, 2012; Marusyk et al., 2012; Zhang and Pellman, 2015). Recent studies using next-generation sequencing have revealed that cancer genomes evolve dynamically through different trajectories even within the same tumor (Almendro et al., 2014; Gao et al., 2016; Marusyk et al., 2014; Ross and Markowetz, 2016). In our work, we have demonstrated *in silico* that re-replication can promote genome plasticity, by generating many diverse genotypes within a population. In cells, incorporation of re-replicating regions into the genome could create site-specific copy gains leading to heterogeneous phenotypes, with potentially desired properties. In a context of natural selection, re-replication may offer a great evolutionary advantage in cells that have lost their normal replication controls, by enabling them to dynamically obtain desired phenotypes and adapt to their environment. Future work will allow such mechanisms to be investigated *in vivo*.

### Conclusions

In summary, we have developed the first mathematical model of DNA re-replication, and extensively simulated it for different hypotheses and model parameters across the complete fission yeast genome. Our *in silico* analysis has elucidated the basic principles that govern DNA re-replication, and indicated how these are manifested depending on the level of analysis: although at the single-cell level re-replication is stochastic and any genomic region is susceptible to amplification, genome-wide patterns of amplification at the population level are robust to different hypotheses and model parameters. These observations highlight that heterogeneity and robustness are emerging and non-contradictory characteristics of DNA re-replication. Importantly, by demonstrating the link between DNA re-replication and genome plasticity, our work may have broad implications for better understanding the onset of genomic instability and cancer evolution.

## Methods

### DNA re-replication model and simulations

A complete mathematical description of the model states, transitions and inputs is given in the Supplemental Note 1. The model was implemented using MATLAB R2016b. Monte Carlo simulations were executed on the HPC cluster of ETH Zurich.

### Statistical methods and data analysis

All methods were implemented in the Statistics and Signal Processing Toolboxes of MATLAB 2016b.

#### Denoising of raw CGH data

For the denoising step, we experimented with various methods (moving average filter, linear polynomial filter, a quadratic polynomial filter and a Savitzky-Golay filter) and a variety of parameter values (e.g., span/window size, degree of the fitted polynomial). From all combinations, a quadratic polynomial fit with a span size of 80 units was chosen as the most appropriate, because of its ability to eliminate noise while fitting the shape and preserving the height of the peaks (Fig. 2 – top in black).

#### Peak finding

To locate the peaks, we implemented a simple peak finding method, which identifies as a peak all local maxima, i.e., all locations where the gradient of the signal changes sign. We also applied an intensity cut-off threshold and set it to 1 to eliminate the peaks whose intensity was below the genome mean. At the same time, a peak-matching step with a threshold of 40 kb was applied, so that local maxima in experimental and simulated data with a linear distance less than 40 kb were assigned to the same peak location. The algorithm identified 12 peaks on Chromosome I, 11 peaks on Chromosome II and 6 peaks on Chromosome III in the denoised experimental data, marked as dotted vertical lines in Fig. 2 (Supplemental Table S2). These numbers were fairly robust with respect to the denoising method but depended largely on the span size and intensity of the cut-off value. The same process was followed for the simulated data and resulted in the identification of 11 peaks in Chromosome I, 5 peaks in Chromosome II and 6 in Chromosome III.

#### Analysis of peak overlap

To assess the overlap between peaks identified in experimental and simulated data, we repeated the same peak matching process using a null model; instead of the simulated peaks, we used 22 randomly-picked genome coordinates and calculated how many coincide with the peaks found in the experimental data using the same window of 40 kb. The procedure was independently repeated 100,000 times; the median overlap score across all repetitions was 2 out of 22 and the maximal overlap score was 10 out of 22, which occurred only once in all 100,000 repetitions.

#### Kernel density estimation

The distribution plots were derived using a kernel density estimate based on a normal kernel and evaluated at 100 equally spaced points.

#### Shannon entropy

To estimate the value of Shannon entropy *H* we first discretized the data to 0 and 1, where 0 corresponds to copy numbers below the genome mean and 1 to simulations above the genome mean. Then *H* is the entropy of a Bernoulli process with probability *p* of two possible and mutually exclusive outcomes, and is defined as follows:

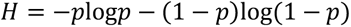

where Pr(*X* = 1) = *p* and Pr(*X* = 0) = 1 − *p*. Entropy will take its maximal value of 1 when *p* = 0.5, i.e., in the case that an origin is amplified in half of the simulations.

#### Principal Component Analysis

To compute the principal components of the data we used the MATLAB function implementation of PCA and visualized the results (variable loadings and principal components) using a biplot.

#### Clustering

To identify groups of similarly amplified profiles in the simulated data, we performed a clustering step using the *k*-means algorithm with a squared Euclidean distance metric. We used the Gap statistic to identify the optimal *k*, a goodness-of-clustering approach that compares the change in within-cluster dispersion with that expected from a reference null distribution (Tibshirani et al., 2001). We estimated the Gap statistic for up to *k*=50 clusters using 100 reference data sets and selected as the optimal the smallest value of *k* for which the value of the Gap statistic is not more than 1 standard error away from the first local maximum. To estimate the stability of the clustering we used the Adjusted Rand Index (ARI), a pairwise metric of similarity between two different clustering assignments. The simple Rand Index (RI) is defined as the number of agreements over all pairs of samples between two different clustering assignments, divided by the total number of pairs, and reflects the probability that a randomly picked pair of samples is consistently found belonging to the same cluster. The ARI is an extension of the RI that additionally corrects for chance (Hubert and Arabie, 1985). To compute the ARI, we ran 100 repetitions of the clustering for the optimal *k* identified as described above, where each repetition was an independent run with a random centroid initialization.

### Experimental methods

#### Strain construction and cell growth

For the construction of the re-replicating strain with the *lys1*+ tag the strain C2566 (h-, LacO::lys1+, LacI-GFP::his7+, ChrI 1.5Mb::TetO-hphMX, Z locus::TetR-tdTomato-natMX, leu1-32, ura4-D18, ade6-M210) kindly provided by Christian Häring was employed (Petrova et al., 2013). To promote re-replication C2566 was crossed with a strain carrying a truncated form of *cdc18* under the nmt1 promoter (*h-, nmt1-d55P6 (leu1+), ura4-D18, leu1-32*) (Baum, 1998). The resulting strain undergoes medium level of re-replication, which is adequate for the manifestation of high cellular heterogeneity but does not severely decrease the viability of the cells. Moreover, medium level of re-replication does not affect the integrity of the nucleus, thus it is possible to detect and quantify Hoechst staining through imaging. This strain was used for the quantification of the *lys1+* locus.

To induce re-replication, cells were grown at 25oC in EMM supplemented with uracil and adenine in the presence of thiamine (5 ug/ml) up to an OD600~0.5. At this point cells were harvested, washed with EMM and diluted to an OD600~0.01 in EMM supplemented with uracil and adenine, both with and without thiamine, and grown at 25oC for 30 hours. Cells were harvested after 30 hours, fixed with 4% PFA for 5 minutes, washed with water and stained with Hoechst for 5 minutes.

#### Cell imaging and image analysis

Cell images were obtained in an Olympus IX83 widefield microscope equipped with a 100x lens (NA 1.46) and a LED light-source. Z-stacks were acquired at a step size of 0.4 μm. Pre-processing, segmentation and signal quantification for GFP and Hoechst were conducted with ImageJ. The top-hat algorithm was applied for background correction. Signals for each channel individually were detected by manual thresholding.

### Data Access

All data generated or analyzed are included in this published article and its supplementary information files. The source code of the model is available here: https://github.com/rapsoman/DNA_Rereplication under an MIT open-source license.

## Acknowledgements

We acknowledge the contribution of Dr. Konstantinos Koutroumpas in the design and implementation of the mathematical model. We thank the Advanced Light Microscopy Facility of the University of Patras. This work was supported by the European Research Council [ERC-StG 281851 and ERC-PoC 755284]; by the State Scholarships Foundation of Greece [short-term fellowship to M.A.R.; PhD fellowship to S.M.] and by the project “Bioimaging-GR” (MIS 5002755), implemented under the Action “Reinforcement of the Research and Innovation Infrastructure”, funded by the Operational Programme "Competitiveness, Entrepreneurship and Innovation" (NSRF 2014-2020) and co-financed by Greece and the EU. Funding for open access charge: Bioimaging-GR.

## Author’s contributions

JL and ZL conceived and supervised the study. JL and ZL designed the mathematical model. MAR implemented the model, ran all simulations, collected and analysed in silico data. MAR, JL and ZL interpreted in silico data. SM, MRG, PN, NNG, ST and ZL designed, performed and/or analysed biological experiments. MAR, JL and ZL wrote the manuscript, with input from all authors. All authors read and approved the final manuscript.

