## Supplementary Figures and Methods for "*In silico* analysis of DNA re-replication across a complete genome reveals cell-to-cell heterogeneity and genome plasticity"

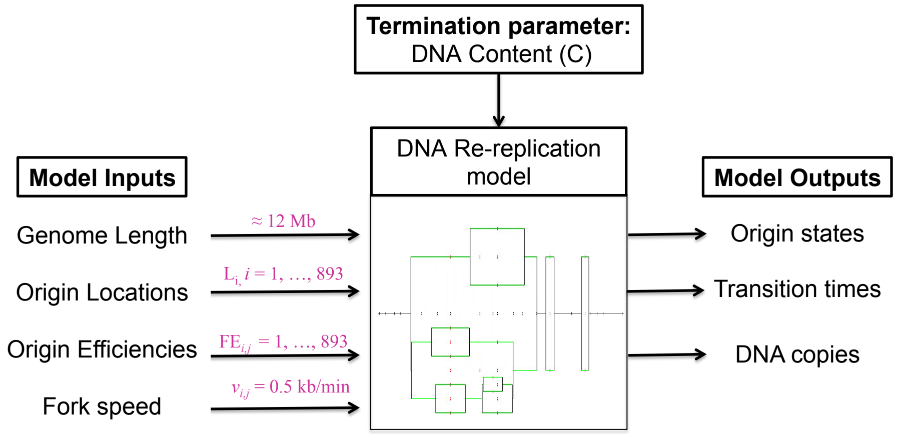


Figure S1: **Model inputs and outputs of the DNA re-replication model.** Model inputs specific for the case of fission yeast reported in purple.

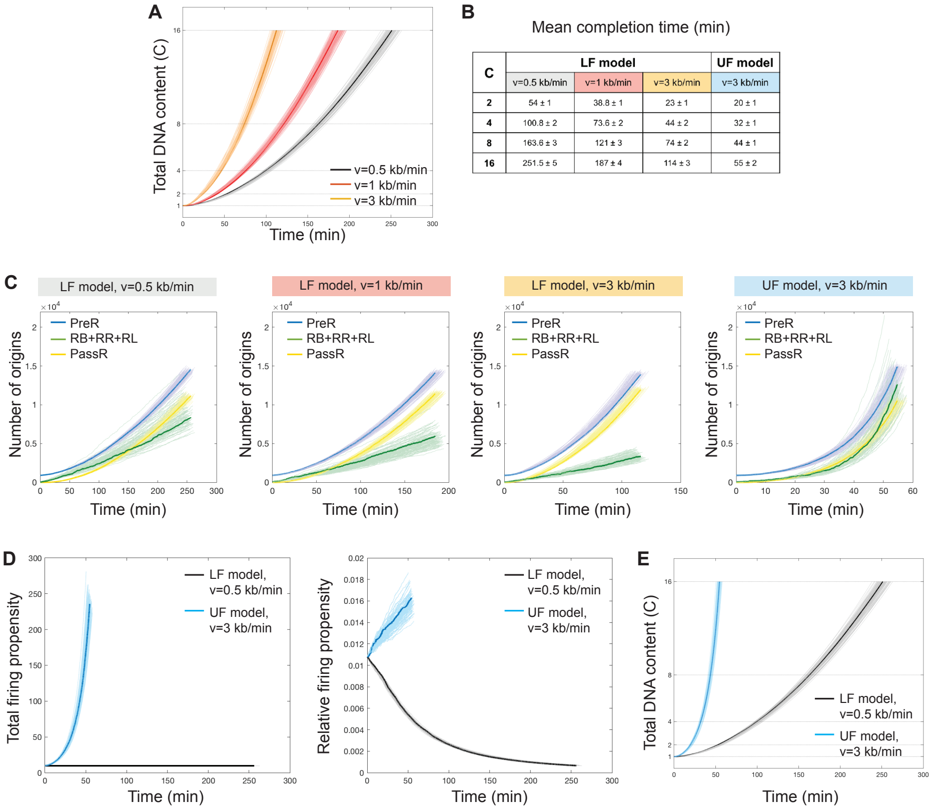


| C | LF model | | | UF model |
|---|---|---|---|---|
| | v=0.5 kb/min | v=1 kb/min | v=3 kb/min | v=3 kb/min |
| **2** | 54 ± 1 | 38.8 ± 1 | 23 ± 1 | 20 ± 1 |
| **4** | 100.8 ± 2 | 73.6 ± 2 | 44 ± 2 | 32 ± 1 |
| **8** | 163.6 ± 3 | 121 ± 3 | 74 ± 2 | 44 ± 1 |
| **16** | 251.5 ± 5 | 187 ± 4 | 114 ± 3 | 55 ± 2 |

Figure S2: **DNA re-replication kinetics for different parameter values and model variations.** Results of 100 Monte Carlo simulations. Different curves correspond to different simulations, and the highlighted curve corresponds to the simulation closest to the mean. Base-case model indicated in black - grey. **(A)** Total DNA content over time for different values of the fork speed. **(B)** Respective completion times in minutes (mean ± standard deviation). **(C)** Total number of origins per state over time; model variations and parameter combinations indicated in the subtitles. Different colors correspond to different origin states. **(D)** Total firing propensity and relative firing propensity (total firing propensity / number of pre-replicative origins) for both model variations. **(E)** Same as in (A), but for the UF and LF model variations.

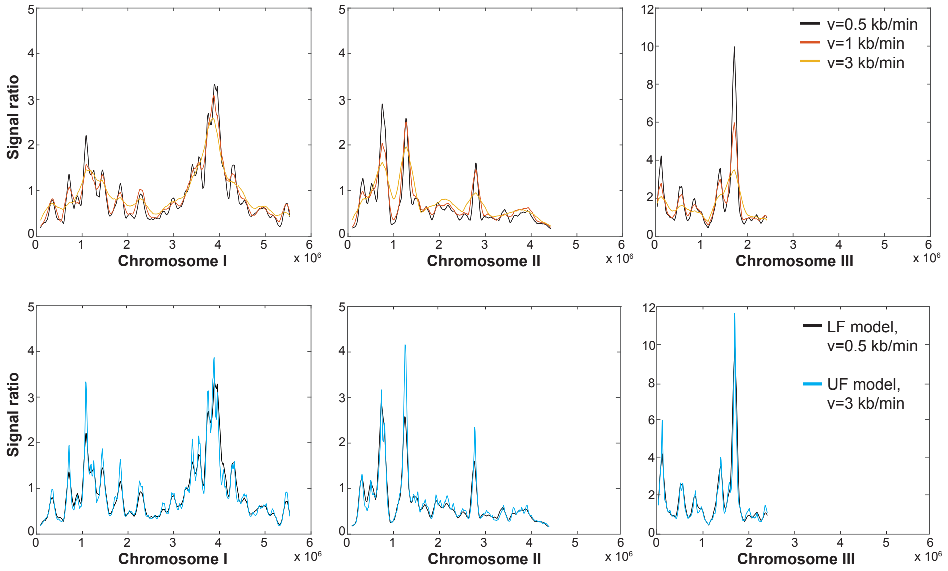


Figure S3: **Amplification levels across the genome for different parameter values and model variations.** Values shown correspond to signal ratios (number of copies / genome mean). Top row: Simulated profiles along the three chromosomes of the fission yeast genome for different values of the fork speed. Bottom row: Comparison of the base-case model (LF model variant, fork speed = 0.5 kb/min) with the best-matching UF model variant.

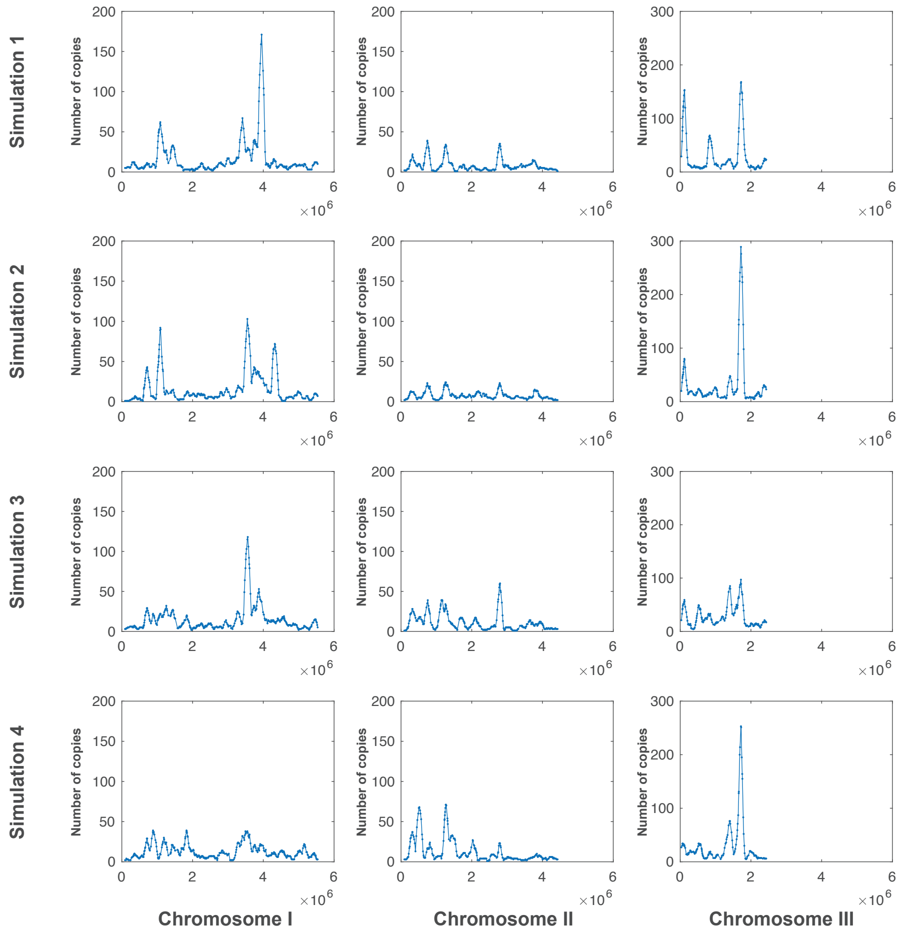


Figure S4: **Individual simulations of the stochastic model lead to markedly different profiles at a single-cell level.** Shown here are number of copies for all origins (marked in circles) across the whole genome, resulting from four random simulations of the model at a total DNA level of $16C$.

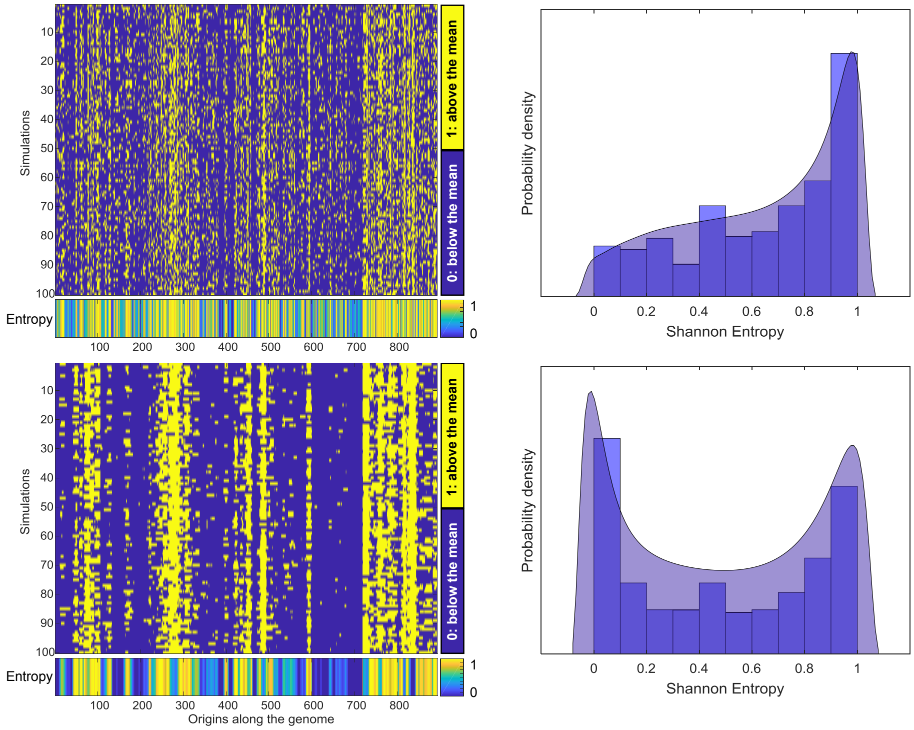


Figure S5: **Quantification of variability using the Shannon entropy index.** Left: heatmaps of discretized data at $2C$ (top) and $16C$ (bottom) with the respective Shannon entropy estimate, shown for all origins across the genome. Right: Distribution of Shannon entropy of all origins for $2C$ (top) and $16C$ (bottom).

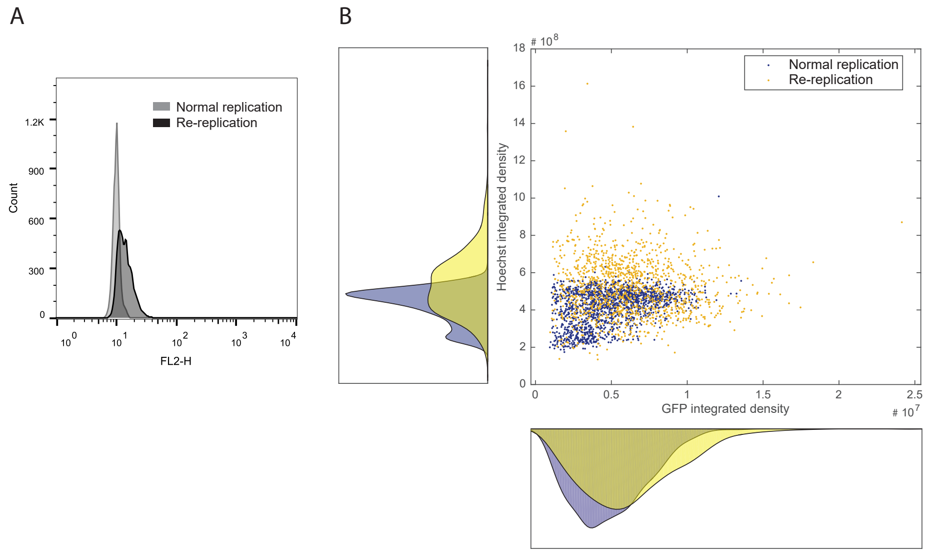


Figure S6: **(A) Overexpression of the d55P6-cdc18 form leads to medium re-replication levels.** Overexpression of the d55P6-cdc18 was induced for 21 hours by thiamine removal (re-replication), or not as a control (normal replication) and the DNA content was monitored using PI (Propidium Iodine) staining and FACS analysis in both conditions. Intermediate re-replication levels were observed upon d55P6-cdc18 overexpression. **(B)** Scatter plot of GFP vs. Hoechst integrated density for normal (blue) and re-replicating (orange) cells, together with their corresponding distributions.

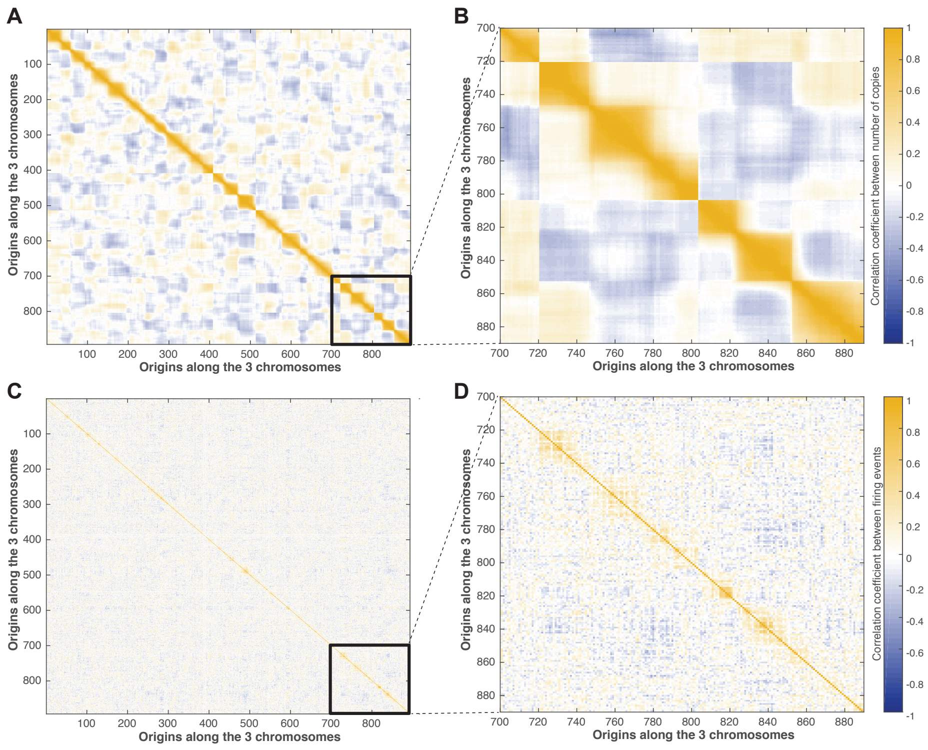


Figure S7: **Correlation coefficients across copy number levels and number of fires. (A-B)** Heatmap of correlations of copy number levels between different origins, shown here for the whole genome (A) and a zoomed in region in the end of Chromosome III (B). Color indicates Spearman correlation coefficients. **(C-D)** Heatmap of correlations of number of firing events between different origins, shown here for the whole genome (C) and a zoomed in region in the end of Chromosome III (D).

### Supplementary Note 1 Model Description

#### Supplementary Note 1.1 Model states and dynamics

Let $L_i, i = 1, , n$ denote the genomic location of each origin and $S_j, j = 1, , m$ denote the DNA strand it is located on. Each origin copy, whether ancestral or offspring, can be identified by its location and strand index and is denoted as $\mathrm{Ori}_{i,j}$. Initially, all origins are located on the parental strand, $j = 1$.

The discrete dynamics of the model are associated with the discrete origin states. Let $q_{i,j}(t) \in \{\text{PreR,RB,RL,RR,PassR,PostR}\}$ denote the state of $\mathrm{Ori}_{i,j}$ at time $t$. It is:

1. **PreR**: Pre-replicative state, if the origin has not yet fired or been passively replicated.

2. **RB**: If the origin has fired, and the replication forks it has initiated in the process are still active in both directions.

3. **RR/RL**: The left/right replication fork initiated by the firing of the origin has met a replication fork from a different origin moving in the opposite direction. The genome between the two origins has been completely replicated on this particular strand, and only the right/left fork initiated by the origin in question is still active.

4. **PassR**: If the origin has been passively replicated by a replication fork initiated by an adjacent origin on the same strand.

5. **PostR**: Both the right and left replication forks initiated by the firing of the origin have met forks initiated by adjacent origins moving in the opposite direction.

Contrary to the normal replication model, the total number of origins in re-replication increases over time, starting with $n$ individual origins on strand $j = 1$. The continuous dynamics model the movement of the replication forks. The continuous dynamics are deterministic and captured by two differential equations whose states represent the number of bases replicated as the forks emanating from an active origin move to the left and right. Let $RF_{i,j}(t), RF_{i,j}(t)$ denote the position of the right and left fork of $\mathrm{Ori}_{i,j}$ at time $t$ and $v$ represent the replication fork speed, measured in bases per minute (see Fig.S8).

It is:

$$\frac{d}{dt}RF_{i,j}(t) = \begin{cases} v(RF_{i,j}(t)) & q_{i,j}(t) \in \{RB, RR\} \\ 0, & \text{otherwise} \end{cases} \tag{S1}$$

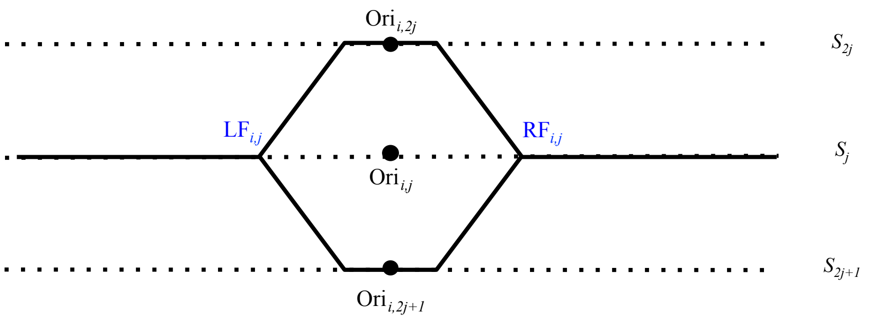


Figure S8: **Indices of origin and fork locations.** Black dots represent origins on different strands, solid lines represent synthesized DNA and dashed horizontal lines different strands. Origin indices are given in black and fork indices in blue.

$$\frac{d}{dt}LF_{i,j}(t) = \begin{cases} -v(RF_{i,j}(t)) & q_{i,j}(t) \in \{RB, RL\} \\ 0, & \text{otherwise} \end{cases} \tag{S2}$$

Uncertainty plays a vital role in re-replication, and it is represented by modeling the time and location of origin firing and re-firing as stochastic events. This is achieved by associating each origin with an intrinsic probability to fire, also referred to as firing propensity, derived from experimentally defined firing efficiencies. Let $FE_{i,j}$ denote the experimentally determined firing efficiency (i.e. the percentage of cells in which an origin fires during one cell cycle) and $TF_{i,j}$ the firing time. The probability that Ori$_{i,j}$ has fired by time $t$ is:

$$P[TF_{i,j} \geq t] = e^{-\lambda_{i,j} t}$$

where $\lambda_{i,j}$ denotes the firing propensity of Ori$_{i,j}$, i.e. the probability to fire in a unit time. It is:

$$\lambda_{i,j} = -\frac{1}{T_c}\ln(1 - FE_{i,j}) \tag{S3}$$

where parameter $T_c$ denotes the completion time of a normal S phase replication. Using the above equation, the firing propensities $\lambda_{i,j}$ can be derived from the experimentally defined firing efficiencies $FE_{i,j}$ in a straightforward manner.

#### Supplementary Note 1.2 Discrete state transitions

Transitions between discrete states, depicted as arrows in Fig.1B, depend on both the continuous and stochastic dynamics of the system. Origin firing is represented through the spontaneous PreR $\rightarrow$ RB transition and is modeled as a stochastic event, governed by an exponentially distributed random variable whose rate depends on each origin's firing propensity. The rest of the discrete state transitions are deterministic and governed by guards, logical statements that depend on the evolution of the continuous state.

The RB$\rightarrow$ RR|RL transitions take place when the left/right replication fork emanating from an active origin meets the right/left replicating fork of the nearest active neighbor to its left/right on the same strand (merging events). Similarly, the RB|RR|RL$\rightarrow$PostR transitions occur when replication forks have merged in both directions. The PreR$\rightarrow$PassR transition occurs when, before an origin has fired, it is passively replicated by a fork emanating from another active origin on the same strand, to the left or right. In contrast to normal DNA replication, origins that have already fired or have been passively replicated can re-fire multiple times. To model this, when origin $\text{Ori}_{i,j}$ fires (transition PreR$\rightarrow$RB) or is passively replicated (transition PreR$\rightarrow$PassR), it generates two origins on two resulting strands (denoted as $\text{Ori}_{i,2j}$ and $\text{Ori}_{i,2j+1}$) which automatically fall back into the PreR state and can thus fire again.

The guard conditions together with the continuous state dynamics allow us to compute transition times for each origin. Let $\text{Ori}_{l,j}, 1 \leq l \leq (i-1)$ denote the closest origin from the left of $\text{Ori}_{i,j}$ on the same strand, for which $q_{l,j} \in \{\text{RB,RR}\}$. Likewise, let $\text{Ori}_{r,j}, (i+1) \leq r \leq n$ denote the closest origin from the right of $\text{Ori}_{i,j}$ on the same strand, for which $q_{r,j} \in \{\text{RB, RL}\}$ (see Fig.S9). In other words:

$$\begin{aligned} \text{Ori}_{l,j} &= \max\{l \leq i \mid q_{l,j} \in \{\text{RB,RR}\}\} \\ \text{Ori}_{r,j} &= \min\{r \geq (i+1) \mid q_{r,j} \in \{\text{RB,RL}\}\} \end{aligned} \quad \text{(S4)}$$

Now let us define the following transitions between the discrete states of the model:

1. **PreR $\rightarrow$ RB**. Occurs when an origin fires and results in the production of 2 offsprings on 2 resulting strands.

2. **RB $\rightarrow$ RR**. Occurs when the left fork of $\text{Ori}_{i,j}$ merges with the right fork of $\text{Ori}_{l,j}$, resulting in $\text{Ori}_{i,j}$ replicating only to the right. For the merging to occur, the following guard condition must be satisfied:

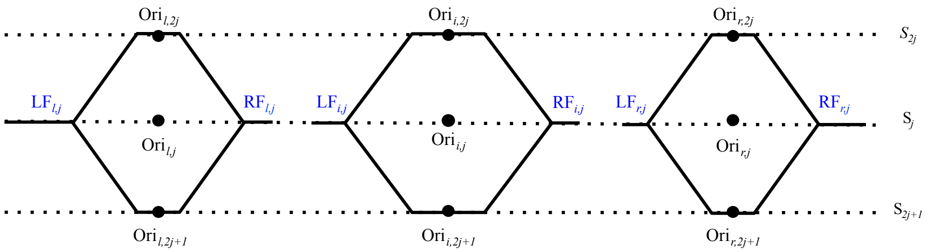


Figure S9: **Indices of origin and fork locations of left and right incoming origins.** Black dots represent different origins, solid lines represent synthesized DNA and dashed horizontal lines different strands. Origin indices are given in black and fork indices in blue.

$$(RF_{l,j} \geq LF_{i,j}) \wedge (RF_{i,j} < LF_{r,j})$$

Then, the time of transition, denoted as $TSL$ (Time to Stop from the Left), is computed as follows:

$$TSL_{i,j} = \frac{L_i - L_l + TF_{i,j} * v + TF_{l,j} * v}{2v}$$

3. **RB** $\rightarrow$ **RL**. Similarly with the above, occurs when the right fork of Ori$_{i,j}$ merges with the left fork of Ori$_{r,j}$, resulting in Ori$_{i,j}$ replicating only to the left. The following guard condition must satisfied:

$$(LF_{r,j} \leq RF_{i,j}) \wedge (LF_{i,j} > RF_{l,j})$$

The time of transition $TSR$ (Time to Stop from the Right) is defined as follows:

$$TSR_{i,j} = \frac{L_r - L_i + TF_{i,j} * v + TF_{r,j} * v}{2v}$$

4. **RR** $\rightarrow$ **PostR:** Occurs when Ori$_{i,j}$ has already merged from the left and subsequently merges from the right. The guard is defined as follows:

$$LF_{r,j} \leq RF_{i,j}$$

The time of transition $TSR$ is defined as above.

5. **RL → PostR:** Occurs when $\text{Ori}_{i,j}$ has already merged from the left and subsequently merges from the right. The guard is defined as follows:

$$RF_{l,j} \leq LF_{i,j}$$

The time of transition $TSL$ is defined again as above.

6. **RB → PostR:** Occurs when $\text{Ori}_{i,j}$ merges simultaneously from the left and the right. The guard is defined as follows:

$$(RF_{l,j} \geq LF_{i,j}) \wedge (LF_{r,j} \leq RF_{i,j})$$

Since the merging events are simultaneous, the time of transition is $TSL = TSR$, computed again as above.

7. **PreR → PassR:** Occurs when $\text{Ori}_{i,j}$ has not fired yet and is passively replicated by any incoming origin from its left or right. Results in the production of two offspring origins, denoted as $\text{Ori}_{i,2j}, \text{Ori}_{i,2j+1}$. For the transition to occur, the following guard statement must be satisfied:

$$(RF_{l,j} \geq L_i) \vee (LF_{r,j} \leq L_i)$$

Then the time of transition (Time of Passive Replication - TPR) is computed as follows:

$$TPR_{i,j} = \begin{cases} \dfrac{L_i - L_l}{v} + TF_{l,j} & \text{if } RF_{l,j} \geq L_i \\ \dfrac{L_r - L_i}{v} + TF_{r,j} & \text{if } LF_{r,j} \leq L_i \end{cases}$$
